# Guardian crypt-base-goblet-cells protect the human colonic stem cell niche by triggering cholinergic calcium signal-dependent MUC2 secretion and luminal flushing

**DOI:** 10.1101/2022.01.04.474646

**Authors:** Nicolas Pelaez-Llaneza, Victoria Jones, Christy Kam, Alvin Lee, Alyson Parris, Sean Tattan, Martin Loader, Jordan Champion, George Russam, Ben Miller, Nathalie Juge, Richard Wharton, Christopher Speakman, Sandeep Kapur, James Hernon, Adam Stearns, Irshad Shaik, Anatu Pal, Alexia Tsigka, Diogenis Batsoulis, Mark Williams

**Affiliations:** School of Biological Sciences, University of East Anglia, Norwich Research Park, Norwich, NR4 7TJ, UK; Quadram Institute, Norwich Research Park, Norwich, NR4 7UQ. UK; Department of General Surgery, Norfolk and Norwich University Hospital, Norwich Research Park, Norwich, NR4 7UY. UK; Department of Histopathology, Norfolk and Norwich University Hospital, Norwich Research Park, Norwich, NR4 7UY. UK

## Abstract

Mucus secreting goblet cells play a vital role in the maintenance of tissue homeostasis. Here we report the discovery of an enigmatic mechanism for the generation of calcium signals that couple cholinergic input to secretion of hydrated mucus in the human colonic stem cell niche. Mechanistic insights for this study were derived from native human colonic crypts and crypt-like organoids expressing MUC2-mNEON. Importantly, we demonstrate that the human colonic stem cell niche is also a cholinergic niche, and that activation of muscarinic receptors initiates calcium signals at the apical pole of intestinal stem cells and neighbouring crypt-base-goblet-cells. The calcium signal ‘trigger zone’ is defined by a microdomain of juxtaposed calcium stores expressing TPC1 and InsP3R3 calcium channels. Co-activation of TPC1 and InsP3R3 is required for generation of cholinergic calcium signals and downstream secretion of hydrated mucus, which culminates in the flushing of the colonic stem cell niche.

## INTRODUCTION

The mucosal surfaces in our body fulfil a diverse array of physiological functions under challenging conditions. A notable example is the lining of the human gut. In addition to facilitating digestion and absorption of nutrients and fluid, the intestinal epithelium forms a vital selective barrier between the mucosal immune system and a barrage of microorganisms, ligands and antigens derived from the hostile gut lumen. Preservation of barrier function is underpinned by rapid stem cell-driven tissue renewal, and by secretion of a protective mucus layer^1^. Compromised barrier function undermines tissue homeostasis and is associated with an increased risk of inflammatory bowel disease and colon cancer^2–4^. Recent advances have provided important insights into the maintenance of epithelial tissue homeostasis and regulation of mucus production by surface goblet cells^2, 5, 6^. However, much less is known about how stem cells and their immediate progenitors function to protect the microenvironment of the stem cell niche from gut luminal contents.

The large intestinal stem cell niche is located at the base of epithelial invaginations called colonic crypts. A population of LGR5^+^ stem cells^7–9^ self-renew on a daily basis^10^ and give rise to progenitor cells that migrate along the crypt-axis and differentiate into absorptive colonocytes or secretory tuft cells, enteroendocrine cells, or goblet cells^11, 12^. The intestinal goblet cell population has recently been shown to be heterogenous with diverse functional features^6, 11^. Sentinel and intercryptal goblet cell subtypes are strategically placed at the crypt opening and within the surface epithelium, respectively^6, 13^. Together with crypt-resident goblet cells, they contribute to the formation of two mucus layers comprising an adherent inner layer that is devoid of bacteria and an outer loose layer^5^, which protect the underlying surface epithelial cells^6^.

The architecture of the colonic crypt adds another dimension to stem cell security by providing a safe harbour for stem cells at the crypt-base, where they are distanced from harmful microbial metabolites and luminal mutagens^14, 15^. Approximately 30-40% of cells that form the axis between the crypt-base and surface epithelium are goblet cells^8^. In the upper region of the crypt, goblet cells are more differentiated, whilst those in the lower region of the crypt still hold proliferative potential, intermingle with stem cells at the crypt-base, and secrete antimicrobial peptides^6, 8, 11^. Mucus secretion by cryptal goblet cells is accompanied by fluid secretion to generate hydrated mucus that flushes the crypt lumen of its contents to maintain a sterile microenvironment^16, 17^. These observations beg the question as to how the interrelated physiological processes of mucus and fluid secretion are functionally coupled in the stem cell niche.

An integral component of the gastrointestinal stem cell niche is the enteric nervous system^18-20^. Cholinergic input along the crypt-axis has been implicated in the regulation of both mucus and fluid secretion^16, 21^, but the mechanism of excitation-secretion coupling is not understood. Previous work in our laboratory has demonstrated initiation of cholinergic calcium signals at the base of human colonic crypts^9, 22^ but the cellular origins, molecular basis and role in excitation-secretion coupling in the intestinal stem cell niche are not known. Here we utilise native human colonic crypts and MUC2::mNEON crypt-like organoids^23^ to address this issue. TPC1 and InsP3R3 cooperation is required for the generation of cholinergic calcium signals in neighbouring colonic stem cells and crypt-base-goblet cells, and also for calcium-dependent luminal secretion of hydrated mucus that flushes the contents of the stem cell niche.

## RESULTS

### A cholinergic niche confers calcium signals upon the human colonic stem cell niche

To further explore the concept of a cholinergic niche^20^ at the base of human colonic crypts^9, 22^ frozen tissue sections of the healthy human colonic mucosa were subjected to immunofluorescence (IF) labelling with cell type-specific markers (Fig. 1 and supplementary Fig. 1)^8^. The intestinal stem cell (ISC) marker OLFM4 labelled slender cells at the crypt-base that were sandwiched between a comparable number of plump MUC2^+^ goblet cells (GCs; Fig. 1a,d and Supplementary Fig. 1a,b,c). GCs in the stem cell niche were also positive for WDCF2 (Fig. 1b and Supplementary Fig. 1b) which has recently been shown to inhibit bacterial growth^11^. Cholinergic tuft cells, which can synthesise acetylcholine^20^, were visualised by double IF labelling for choline acetyltransferase (ChAT) and advillin^24^, and were predominantly distributed in the lower-half of the crypt-axis (Supplementary Fig. 1d). Cholinergic tuft cells were relatively rare compared to ISCs and GCs (c.f. Figs 1d&e), as were Chromogranin A^+^ enteroendocrine cells (Fig. 1g). ChAT expression in the mucosa and isolated epithelial crypts was confirmed by RT-PCR (Supplementary Fig. 1e). The close proximity of cholinergic neurons to the human colonic stem cell niche was established by 3D rendering a series of confocal images taken through the colonic mucosa subjected to IF labelling for the neuronal marker beta tubulin III and ChAT. (Fig. 1f). The presence of neuronal and non-neuronal sources of acetylcholine in the stem cell niche prompted an analysis of epithelial cell subtypes that express muscarinic receptors. M3AChR mRNA expression in patient-matched mucosa, isolated crypts and organoids was confirmed by RT-PCR (Fig. 1h). M3AChR IF Iabelling of the native mucosa predominated at the crypt base (Fig. 1g)^9^ and IF labelling of naïve isolated colonic crypts and crypt-like organoids unveiled M3AChR expression by LGR5^+^ ISCs and MUC2^+^ GCs (c.f. relatively low levels in CGA^+^ enteroendocrine cells and ChAT^+^ tuft cells; Fig. 1g and Supplementary Fig. 1g). Expression of M1AChR and M5AChR at the colonic crypt-base (Supplementary Figs 1f,h,i) reinforces the concept that the human colonic stem cell niche is also a cholinergic niche (Fig. 1i).

**Figure 1.**
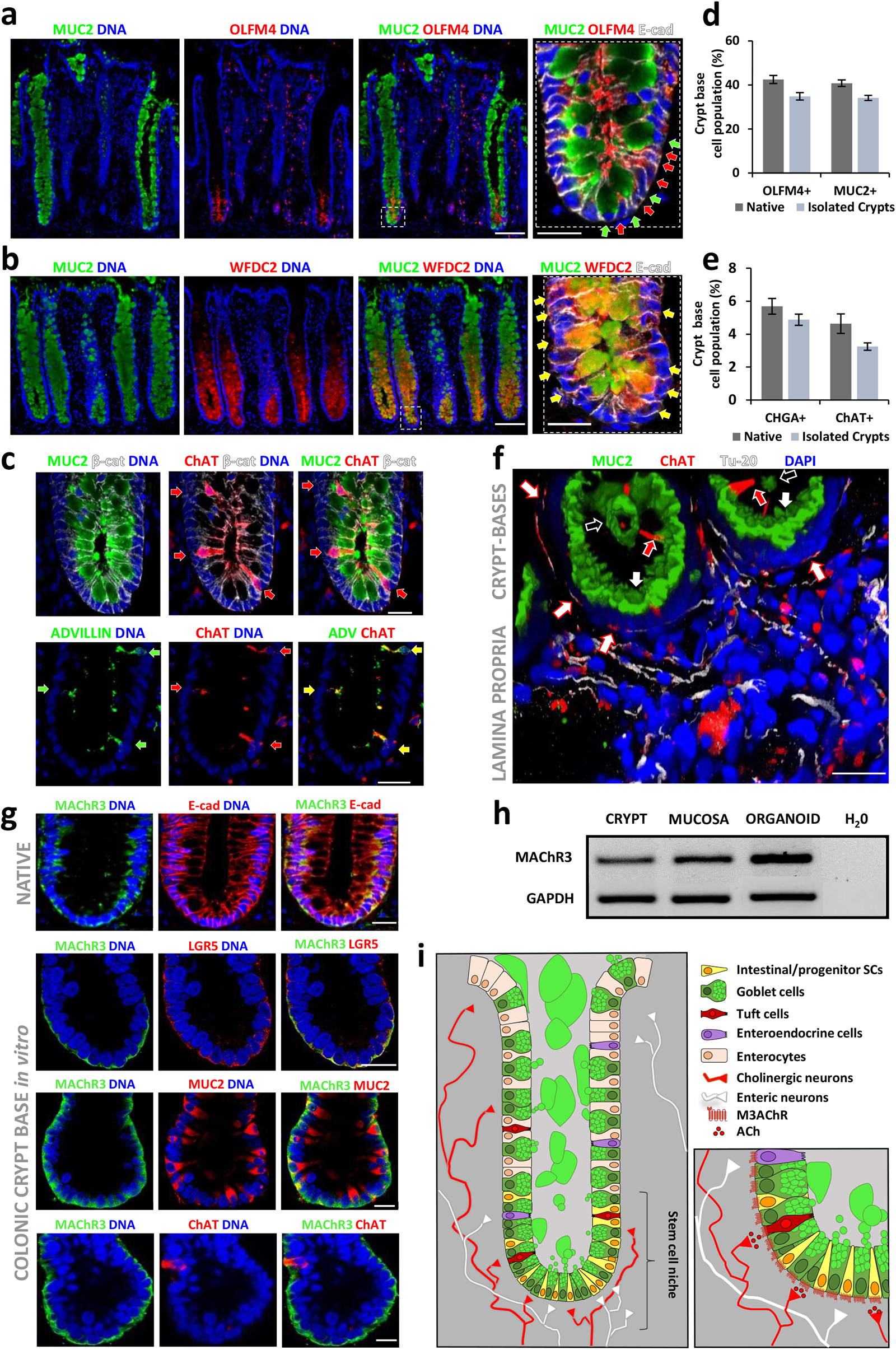
The stem cell niche at the human colonic crypt base is also a cholinergic niche. (a) Cellular composition of the intestinal stem cell niche at the base of human colonic crypts in vivo; immunofluorescent labelling of interdigitated MUC2 (green)-positive goblet cells and OLFM4 (red)-positive stem cells at the base of crypts in the native human colon. (b) Co-labelling (yellow arrowheads) of MUC2 (green)- and WFDC2 (red)-positive goblet cells at the base of native human colonic crypts; scale bars = 50μm/25μm. (c) Double immunolabelling of ChAT (red) with MUC2 (green, top) and the tuft cell marker advillin (green, bottom) in cryosections of the human colonic mucosa, scale bars 25μm; co-labelling indicated by yellow arrowheads. Quantification of abundant (d) and rare (e) cell types in the native and isolated human colonic crypt stem cell niche. (f) A 3D render produced from a confocal series of immunofluorescent images of the human colonic submucosa; MUC2 immunolabelling visualises goblet cells (white arrowheads) and luminal mucus at the colonic crypt-base (open arrowheads), ChAT labels epithelial tuft cells within the crypt (white outlined red arrows) and cholinergic neurons that are double-labelled with the neuronal specific b-tubulin III antibody, TU-20 (red open arrows); scale bar = 100μm. (g) Immunofluorescent labelling of human colonic epithelial M3 muscarinic receptors and cell type markers in the native mucosa and cultured colonic crypt-base; intestinal stem/progenitor cells – LGR5, goblet cells – MUC2, tuft cells – ChAT; scale bar = 20μm. (h) RT-PCR of M3 muscarinic receptor in native mucosa, isolated crypts and cultured organoids. (i) Schematic representation of the large intestinal cholinergic-stem cell niche. Immunolabelling patterns are representative of tissue sections and isolated crypts derived from N>4 different subjects. Unpaired t tests were undergone for (d), (e) denoting no significant differences.

We have previously demonstrated that acetylcholine initiates calcium signals in the colonic stem cell niche via activation of M3AChRs followed by signal propagation along the topological axis of the tissue; this is confirmed here for the acetylcholine analogue Carbachol (CCh, 10 μM, approximate EC50; Fig. 2a,b and Supplementary Fig. 2a,b; Supplementary Movie 1)^9, 22, 25^. The action of CCh was mimicked by the muscarinic receptor agonist Oxotremorine and inhibited by the M3AChR-selective antagonist 4-DAMP (Figs 2c-f). Confocal imaging was then used to analyse colonic crypt calcium signalling at the single cell level in situ. Cholinergic calcium signals initiated at the apical pole of slender cells (i.e. of typical stem cell morphology) located within the stem cell niche (Fig. 2g), and spread via an apparent apical path throughout the crypt-base (Fig. 2g,h; Supplementary Movie 2). Post-hoc IF labelling confirmed that cholinergic calcium signals initiated in OLFM4^+^ ISCs before registering in neighbouring cell types, including MUC2^+^ crypt-base-GCs (Fig. 2i,j and Supplementary Fig. 2c), which exhibited a similar relative increase in cytosolic calcium amplitude (Fig. 2k,l).

**Figure 2.**
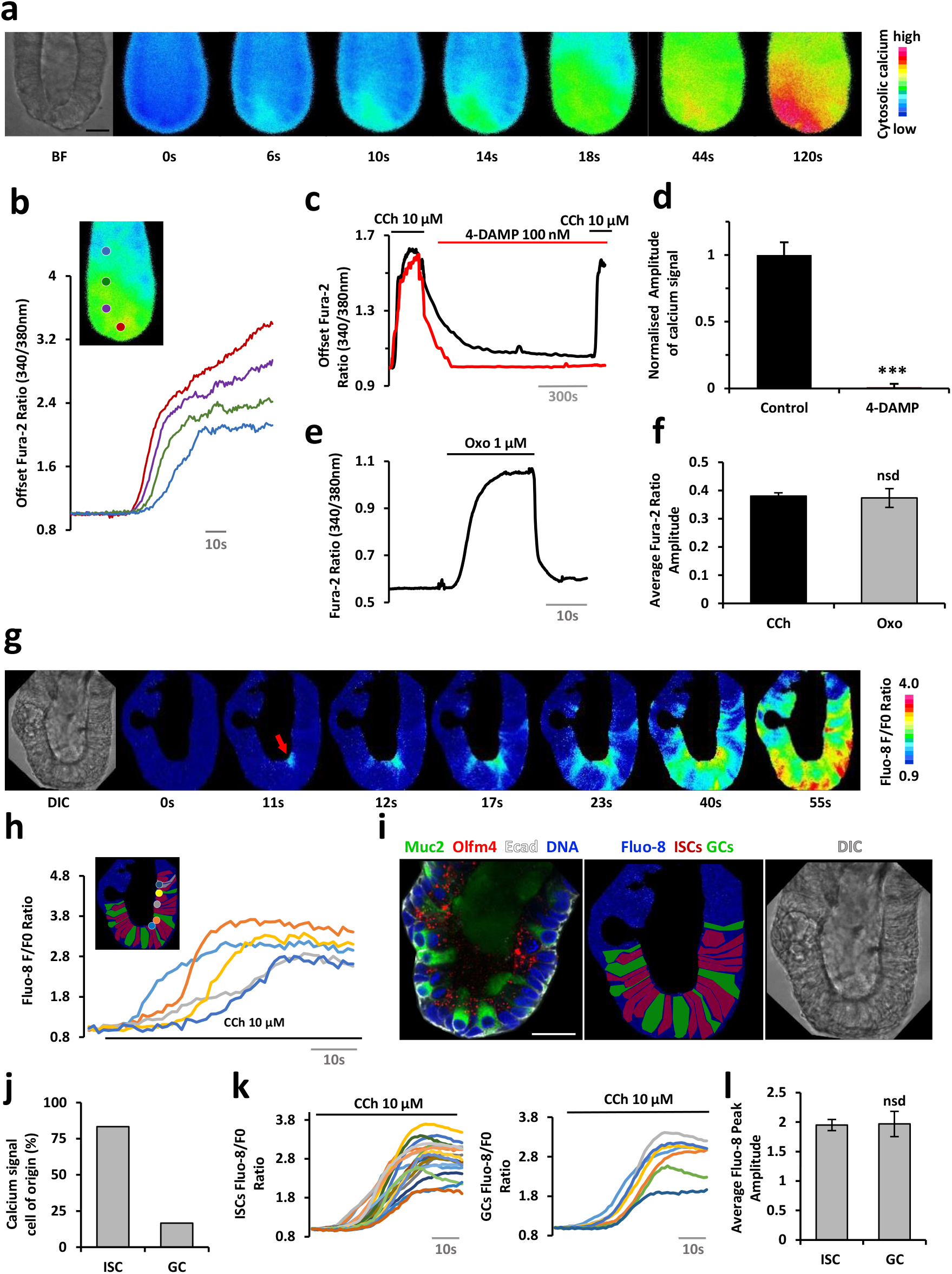
Cholinergic input stimulates calcium signals in intestinal stem cells and goblet cells at the human colonic crypt-base. (a) Spatiotemporal characteristics of colonic crypt calcium signals in the cholinergic stem cell niche; Fura-2 ratio images of a human colonic crypt stimulated with Carbachol (10 µM) and (b) kinetics of cholinergic calcium signal generation in regions of interest placed along the colonic crypt-axis (N>10 subjects, n>50 crypts). Effect of M3AChR-selective inhibitor 4-DAMP (100 nM) on Carbachol-induced calcium signal generation (c) and amplitude (d) at the colonic crypt-base (N>3, n>10). Activation of calcium signals by the muscarinic receptor agonist Oxotremorine (1 µM) (e) with respect to Carbachol (f) (N>3, n>10 crypts). (g) Single cell calcium signals at the colonic crypt base/stem cell niche of an organoid resolved in situ by Fluo-8 confocal fluorescence F/F0 ratio imaging (red arrowhead indicates site of calcium signal initiation). (h) Kinetics of single cell cholinergic calcium signals along the organoid crypt-axis. (i) Post-hoc immunophenotyping of colonic crypt cell types exhibiting Carbachol-stimulated cholinergic calcium signals (goblet cells - MUC2, green; intestinal stem cells - OLFM4, red). (j) Post-hoc identification of calcium signal cell-of-origin and calcium signal kinetics (k) and amplitude (l) in intestinal stem cells (ISCs) and goblet cells (GCs) (N=2, n=6 organoid crypt bases). Scale bars = 25 µm, ***P<0.001. Resting ratio of some Fura-2 ratio traces has been offset to aid comparison of responses in (b&c); there was no significant difference in the resting Fura-2 ratio for different experimental groups in (c). P values calculated via unpaired t-tests.

### Cholinergic calcium signals stimulate MUC2 secretion in the human colonic stem cell niche

Cultured human colonic crypts preserve the tissue topology, cellular polarity and cellular diversity associated with the native human gut epithelium^8^. The confocal (x,y) orientation plane for GC polarity (Fig. 3a) gave a clear visualisation of GC MUC2 granular content following stimulation with CCh; Fig. 3b illustrates a pseudo-time representation of GCs in progressive states of MUC2 granule exocytosis leading to GC MUC2 depletion.

**Figure 3:**
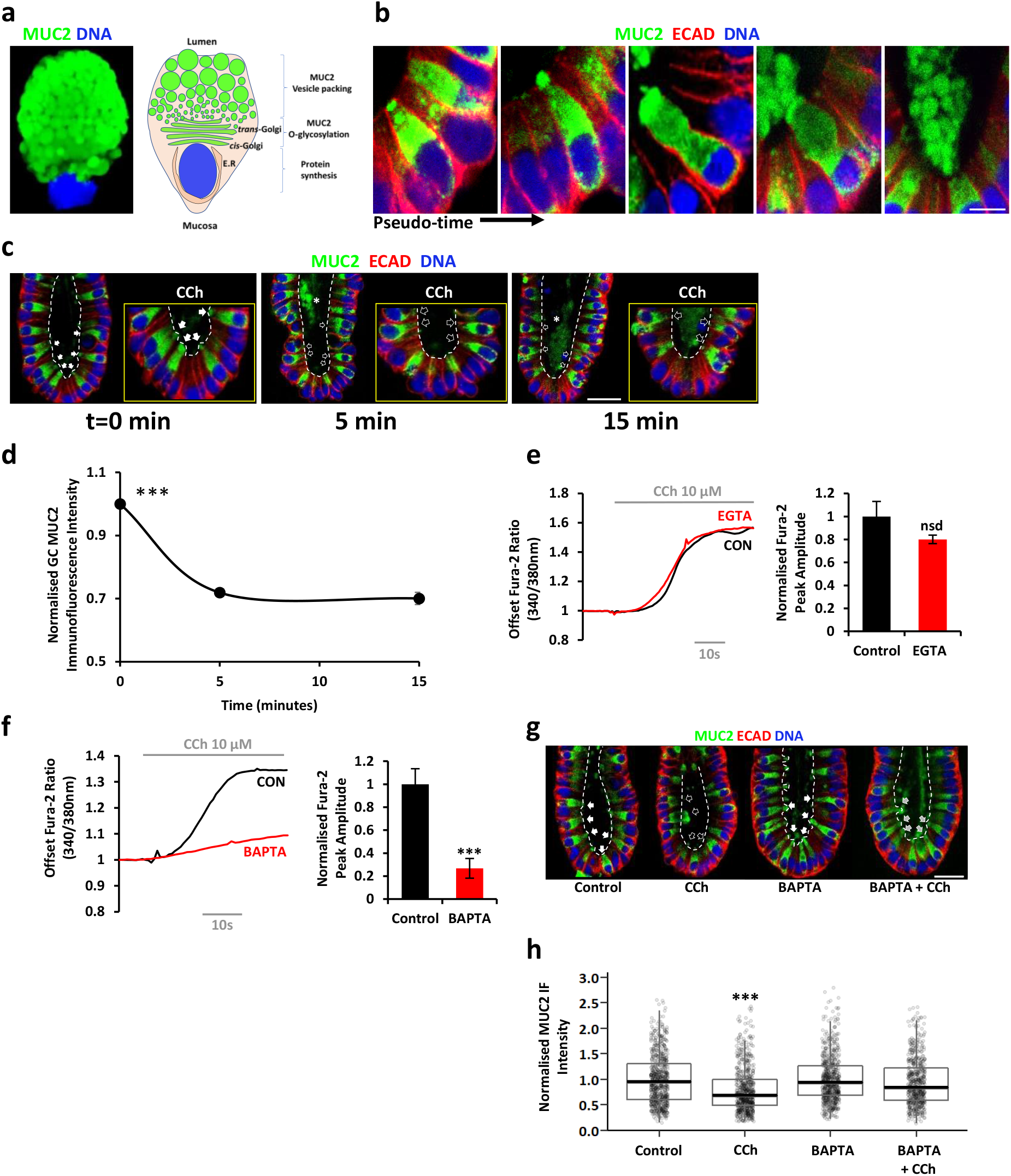
Muscarinergic mobilisation of intracellular calcium is a requirement for mucus secretion at the human colonic crypt base. (a) Fluorescence immunolabelling (MUC2, green) of a single human colonic crypt base goblet cell with an accompanying schematic illustration. (b) Representative examples of the progressive stages of mucus granule expulsion by crypt base goblet cells in situ following stimulation with Carbachol (10 µM, pseudo-time) visualised by post hoc MUC2 immunolabelling (green); scale bar = 10 µm. (c) Absolute time course for Carbachol-stimulated (10 µM, time minutes) depletion of MUC2 from crypt-base goblet cells and (d) corresponding kinetics for goblet cell MUC2 immunofluorescence intensity (normalised to unstimulated control at matched time points); N=3, crypt≥10, GC>100. Scale bar = 25 µm. (e) Carbachol-stimulated calcium mobilisation in the absence of extracellular calcium (Hepes-buffered saline minus calcium, plus EGTA, 1 mM) and calcium signal amplitude analysis; Fura-2 resting ratio offset to 1.0 to aid comparison. (f) Suppression of Carbachol-induced increase of cytoplasmic calcium peak amplitude by pre-incubation with the calcium chelator BAPTA-AM (66 mM, 45 min.), Fura-2 resting ratio offset to 1.0 for clarity. Effects of BAPTA-AM pre-incubation on Carbachol (10 µM, 5 mins)-stimulated mucus depletion from crypt-base goblet cells, (g) example MUC2 immunofluorescence images and (h) image analysis of relative MUC2 goblet cell content. White asterisk - secreted mucus granules, filled arrowheads – goblet cells replete with mucus, empty arrowheads - goblet cells depleted of MUC2, grey filled arrowheads – partially emptied/re-filled MUC2 goblet cells. N=3, crypt≥10, GC>100. Scale bar = 25 µm. ***P<0.001, significantly different to all other groups. No significant difference in resting Fura-2 ratio for different experimental groups in (e) and (f). One-way ANOVA was undergone for (d), an unpaired t test was undergone for (e), a Kruskal-Wallis rank sum test with Dunn post hoc was undergone for (h).

Unstimulated GCs exhibit characteristic apical ‘bulbs’ of MUC2-labelling. The kinetics of GC MUC2 depletion peak at 10-15 min (Figs 3c,d and Supplementary Figs 3a,b). The dependence of GC MUC2 secretion on elevated intracellular calcium levels was examined next. Cholinergic calcium signals were not affected by acute removal of extracellular calcium (Fig. 3e), but could be buffered by pre-incubation with BAPTA-AM (Fig. 3f), which also suppressed CCh-induced GC depletion of MUC2 (Fig. 3g,h). Calcium has been implicated in luminal stimulation of MUC2 secretion by differentiated GCs in the mouse gut^13^ and organoids^17^ but the origins and mechanism for the calcium signal is not yet fully understood.

### Co-operation between endosomal and ER calcium stores is a requirement for cholinergic calcium signal generation in the colonic ISC niche

It is clear that the cholinergic calcium signal originates from an intracellular calcium store (Fig. 3e). The cellular location of endolysosomal and endoplasmic reticular (ER) calcium stores in the stem cell niche was visualised by lysotracker red fluorescence and KDEL IF labelling, respectively. Puncta exhibiting intense Lysotracker fluorescence were prominent at the apical pole and diminished in intensity towards the basal pole of cells in the ISC niche (Fig. 4a). Conversely, IF labelling of the ER marker KDEL was concentrated around the basal nuclei and extended toward the apical pole (Fig. 4a). We next demonstrated the expression of candidate intracellular calcium channels InsP3Rs1-3, RyRs1-3 and TPCs1&2 by RT-PCR analysis of mRNA derived from isolated epithelial crypts (Fig. 4b), the native mucosa and crypt-like colonic organoids (Supplementary Figs 4e,f). IF labelling for InsP3R2, RyR1-3 and TPC2 predominated at the basal pole of crypt-base cells (Fig. 4c,d), whereas InsP3R1 IF labelling extended from the basal pole toward the apical pole (Fig. 4c,d). Strikingly, TPC1 IF co-labelled discrete Rab11^+^ endosomes that were juxtaposed to InsP3R3^+^ puncta situated beneath the cellular apical membrane. These so called ‘microdomains’ were present within cells of slender ISC-like morphology and plump cells of GC-like morphology. TPC2 IF labelling exhibited partial co-localisation at the basal pole with Rab5^+^ puncta (Fig. 4d). The apical and basal location of TPC1 and TPC2, respectively, was confirmed in isolated single GCs (Fig. 4f). Similar expression patterns for intracellular calcium channels were exhibited by crypt-like organoids (Supplementary Fig. 4a-d). Of note, CD38, an enzyme that can modulate NAADP production (a renowned activator of TPCs)^26^, also co-localises with Rab11^+^ puncta at the apical pole of MUC2^+^ GCs and neighbouring slender ISCs (Fig. 4e).

**Figure 4:**
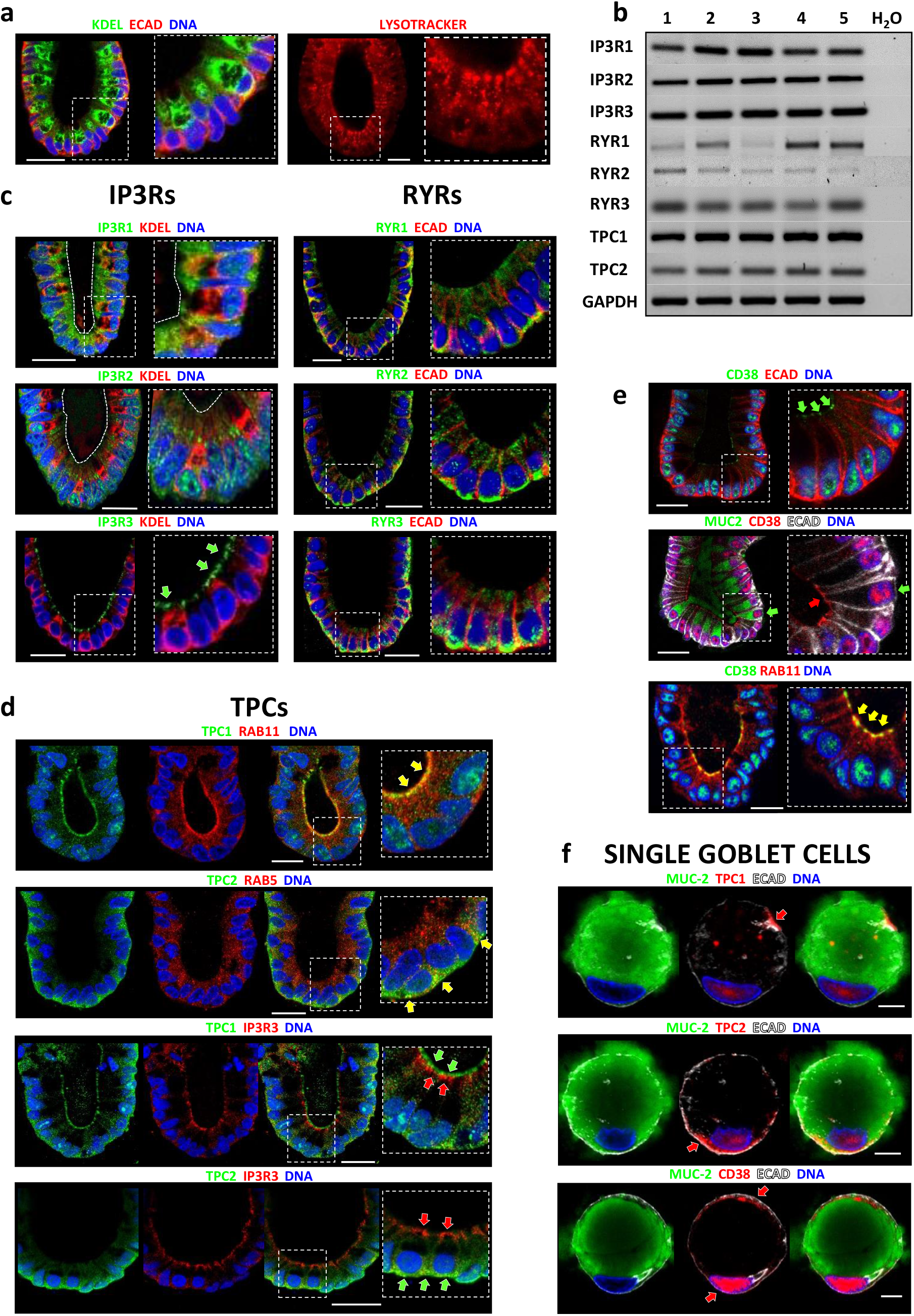
Polarised distribution of intracellular calcium stores and calcium release channels in the stem cell niche at the human colonic crypt-base. (a) Immunolocalisation of the endoplasmic reticulum (KDEL) and fluorescence imaging of endolysosomes (lysotracker, 1mM for 1 hour) in colonic crypt base cells; N≥3 subjects, n≥10 crypts. Scale bar = 25µm. (b) RT-PCR of intracellular calcium release channel mRNA transcript expression in acutely isolated human colonic crypt samples (N=5 subjects). (c) Immunolocalisation of InsP3R1, InsP3R2, InsP3R3, RYR1, RYR2, RYR3, in combination with endoplasmic reticulum marker KDEL and/or E-Cadherin; N ≥ 3, n ≥ 10; Scale bar = 25µm. (d) Immunolocalisation of CD38 in combination with goblet cell marker MUC2 and/or E-cadherin; N≥3, n≥10 crypts. Scale bar = 25µm. (e) Immunolocalisation of TPC1 and TPC2 in combination with recycling (RAB11) or early (RAB5) endolysosomal markers, or InsP3R3; N≥3, n≥10 crypts. Scale bar = 25µm. (f) Immunolocalisation of TPC1, TPC2 and CD38 in combination with MUC2 in single goblet cells; N≥3, n≥10; scale bar = 5µm. (f) Expression of CD38 and TPCs in single goblet cells. Yellow arrowheads indicate co-labelling, and red and green arrowheads indicate distinct immunolabelling of colour-coded proteins.

We next assessed the roles of ER and endolysosomal calcium stores for generation of cholinergic calcium signals in the stem cell niche. Depletion of ER stores by the SERCA pump inhibitor cyclopiazonic acid (CPA), in the absence of extracellular calcium, markedly suppressed the cholinergic calcium response (Fig 5a). A predominant role for mobilisation from ER stores is supported by sensitivity of the cholinergic calcium response to phospholipase C inhibition (U73122^27^, Fig. 5b) and InsP3R inhibition by caffeine^28, 29^ (Fig. 5c), but not by Xestospongin C^30^ nor 2-APB^28^ (Supplementary Fig. 5a,f). The cholinergic calcium response was also diminished by the RyR blockers dantrolene and ryanodine (Fig. 5d)^31^.

**Figure 5:**
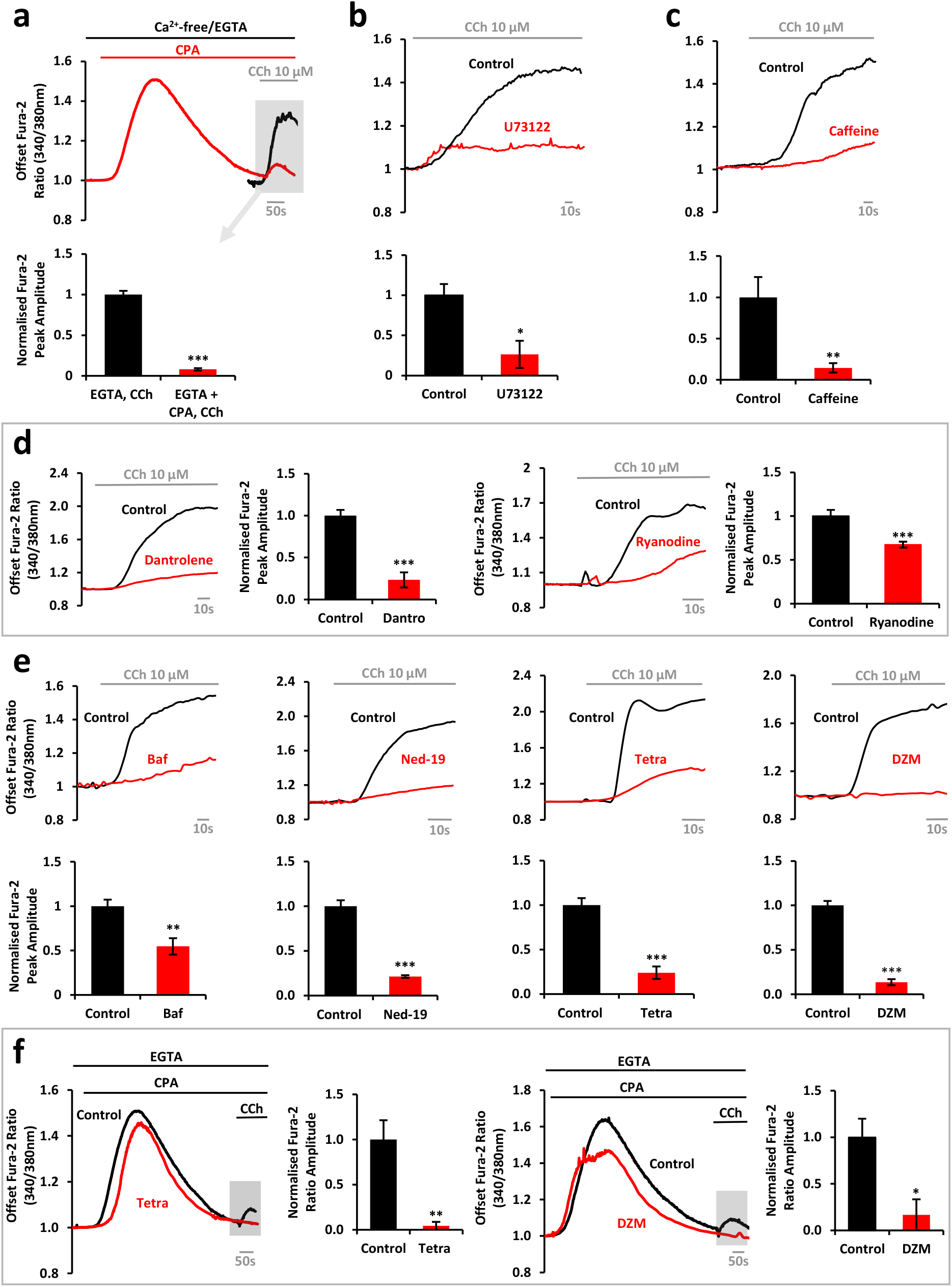
Muscarinergic calcium signals are dependent on cooperation between two pore channels, InsP3 receptors and Ryanodine receptors. (a) Top - Effects of ER calcium store depletion on Carbachol-induced calcium response. Fura-2 ratio responses to Carbachol (10 µM) stimulation following pre-treatment with Ca^2+^-free/EGTA (1 mM) HBS in the presence (red) and absence (black) of SERCA pump inhibitor CPA (20 µM); bottom - bar chart of Carbachol response peak amplitude normalised to that obtained in the absence of CPA. (b) Effects of PLC inhibitor U73122 (10 µM; 30 min.) on Carbachol-induced calcium signal generation, and bar chart of normalised peak amplitude of the responses. (c) Effects of InsP3 receptor inhibitor Caffeine (10 mM, 30 min.) on Carbachol-induced calcium signal generation, and the bar chart of peak amplitude of the responses. (d) Effects of RyR inhibitors Dantrolene (30 µM, 2 hrs) and Ryanodine (50 µM, 30 min.) on CCh-induced calcium signal generation and the corresponding bar chart of peak amplitude of the response. (e) Effects of the lysosomotropic agent Bafilomycin (2.5 µM, 2 hrs) and TPC inhibitors Ned-19 (500 µM, 2 hrs), Tetrandrine (20 µM, 2 hrs) and Diltiazem (250 µM, 30 min.) on Carbachol-induced calcium signal generation and the corresponding bar chart of peak amplitude of the response. (f) Effects of TPC inhibitors Tetrandrine (20 µM, 2 hrs) or Diltiazem (250 µM, 30 min.) on the residual Carbachol-induced calcium response following ER calcium store depletion by the SERCA pump inhibitor CPA (20 µM; HBS minus calcium, plus EGTA, 1 mM), traces and bar charts of peak amplitude. N≥2 subjects, n≥9 crypts in each case. Resting ratio of some Fura-2 ratio traces has been offset to aid comparison of responses in (a-e); there was no significant difference in the resting Fura-2 ratio for different experimental groups in (a-e). ***P<0.001, **P<0.01, and *P<0.05 indicate a statistical difference to all other groups at the indicated level of significance. P values calculated via unpaired t-tests.

Endolysosomes are acidic calcium stores and their calcium load can be dissipated by blocking the proton V-ATPase^32^. The cholinergic calcium response was abrogated by pre-treatment with bafilomycin (Fig. 5e). Calcium release from endolysosomes is commonly mediated by two pore channels, TPC1 and TPC2^32^. TPC channel blockers Ned-19^33^, tetrandrine^34^ and diltiazem^34, 35^ each reduced the amplitude of the cholinergic calcium response (Fig. 5e). TPC channel blockers also abolished the residual response to CCh following CPA-mediated depletion of the ER in the absence of extracellular calcium (see shaded aspects in Figs 5a,f). Calcium responses evoked by TPC agonists were also blocked by TPC inhibitors (Supplementary Fig. 5b).

### Cholinergic calcium signals couple mucus secretion with fluid secretion in the human colonic stem cell niche

Pre-treatment of cultured human colonic crypts with intracellular calcium channel inhibitors caffeine, dantrolene, Ned-19 (Fig. 6a,b,c) and tetrandrine (Supplementary 6b) suppressed CCh-induced MUC2 depletion of GCs, as did inhibition of M3AChRs with 4DAMP (Supplementary Fig. 6c). The InsP3R inhibitor 2-APB, which did not abrogate CCh-induced MUC2 depletion, did not suppress CCh-induced MUC2 depletion (Supplementary Figs 5a & 6a). Having utilised the MUC2 IF depletion assay to forge functional links between calcium mobilisation and mucus secretion, we next generated MUC2::mNEON human colonic crypt-like organoids to investigate the impact of the cholinergic calcium signalling pathway on coupling mucus and fluid secretion to flush the stem cell niche (Supplementary Fig. 7a-d)^23, 36^.

**Figure 6.**
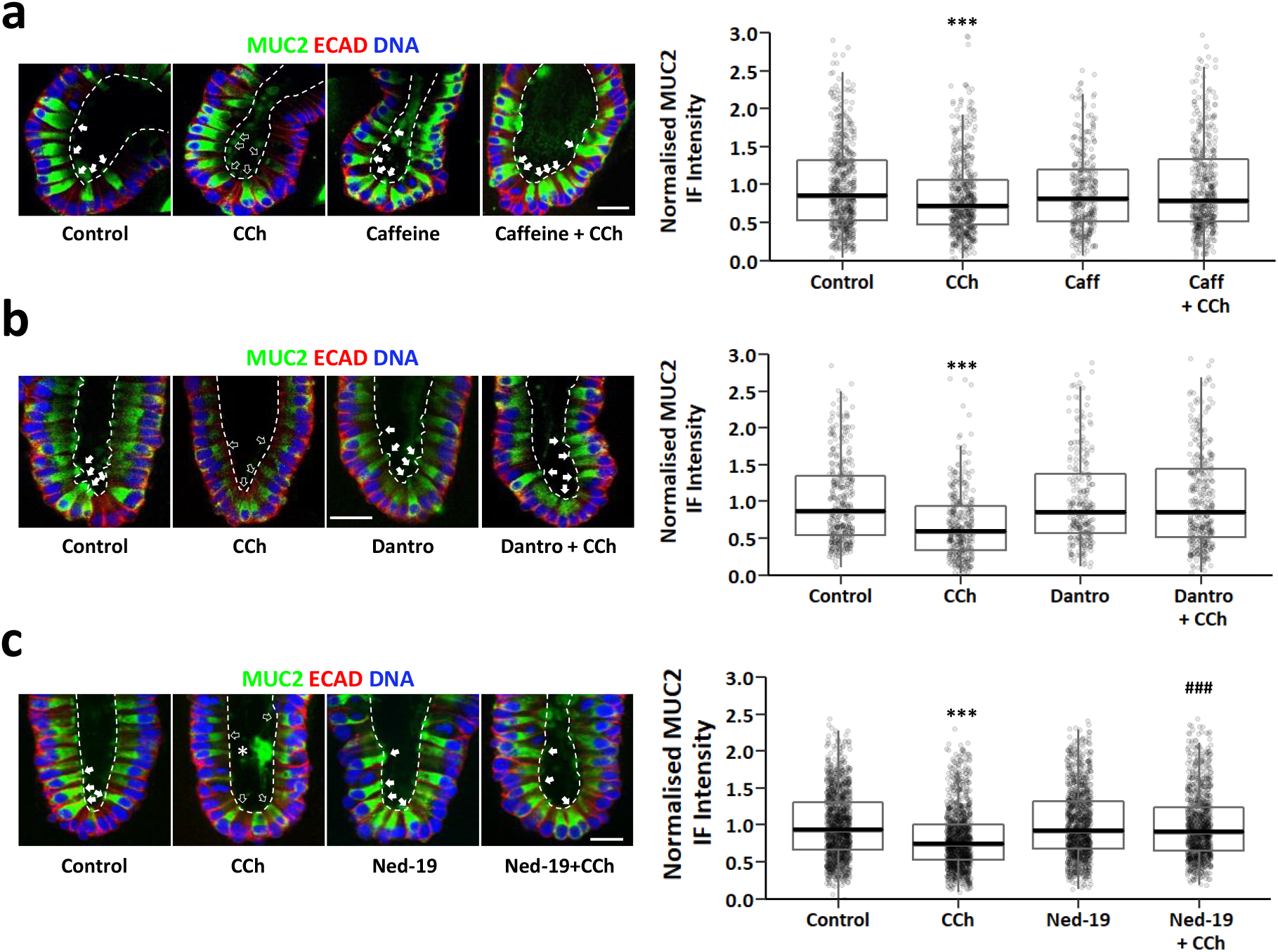
Cholinergic excitation-mucus secretion coupling in the human intestinal stem cell niche is dependent on cooperation between two pore calcium channels, InsP3 receptors and ryanodine receptors. Representative MUC2 (green) immunofluorescence images and summary box plots for image analysis of MUC2 immunofluorescence intensity. Effects of (a) InsP3R inhibitor Caffeine (10mM, 30 min.), (b) RyR inhibitor Dantrolene (30μM, 2 hrs) and (c) TPC inhibitor Ned-19 (500μM, 2 hrs), on Carbachol-stimulated (10µM, 5min) mucus depletion from colonic crypt-base goblet cells are presented. Asterisks on images - secreted mucus granules, empty arrowheads – goblet cells depleted of MUC2, filled arrowheads – goblet cells replete with mucus. N=2 subjects, n>10 crypts, GC>100 in each case, scale bar = 20μm. ***P<0.001 with respect to all other groups; ##P<0.01 with respect to control. P values calculated via Kruskal Wallis rank sum tests with a Dunn post-hoc.

MUC2::mNEON heterozygous crypt-like organoids displayed MUC2-mNEON fluorescence that exhibited a granular appearance at the apical pole and which co-localised with MUC2 immunofluorescence (Fig. Supplementary Fig. 7d). MUC2::mNEON organoid lines also exhibited cholinergic calcium signals with similar spatiotemporal characteristics to wild type organoids (Supplementary Movie 3; Supplementary Fig. 7 e-h); cholinergic calcium signals initiated in slender MUC2-mNEON^-^ cells located at the crypt-base and spread to neighbouring MUC2-mNEON^+^ cells and achieved a similar relative peak amplitude within 10 seconds (Supplementary Fig. 7e-h; Supplementary Movie 3). Cholinergic input also stimulated an immediate increase in luminal MUC2-mNEON fluorescence followed by morphometric changes that were reminiscent of cellular secretion (Fig. 7a-c; Supplementary Movie 4)^9^; the cross-sectional area of the crypt lumen increased in size and the apical-to-basal dimension exhibited a secretory volume decrease (Fig. 7e,f)^9^. An increase in luminal MUC2-mNEON fluorescence (Fig. 7c) was accompanied by a pronounced reduction in MUC2-mNEON ‘theca’ volume (Fig. 7f) consistent with expulsion of MUC2-mNEON into the crypt lumen. Pre-incubation with the TPC inhibitor Ned-19 (Fig. 7g) or the InsP3R inhibitor caffeine (Fig. 7h) suppressed all these cholinergic-induced effects.

**Figure 7:**
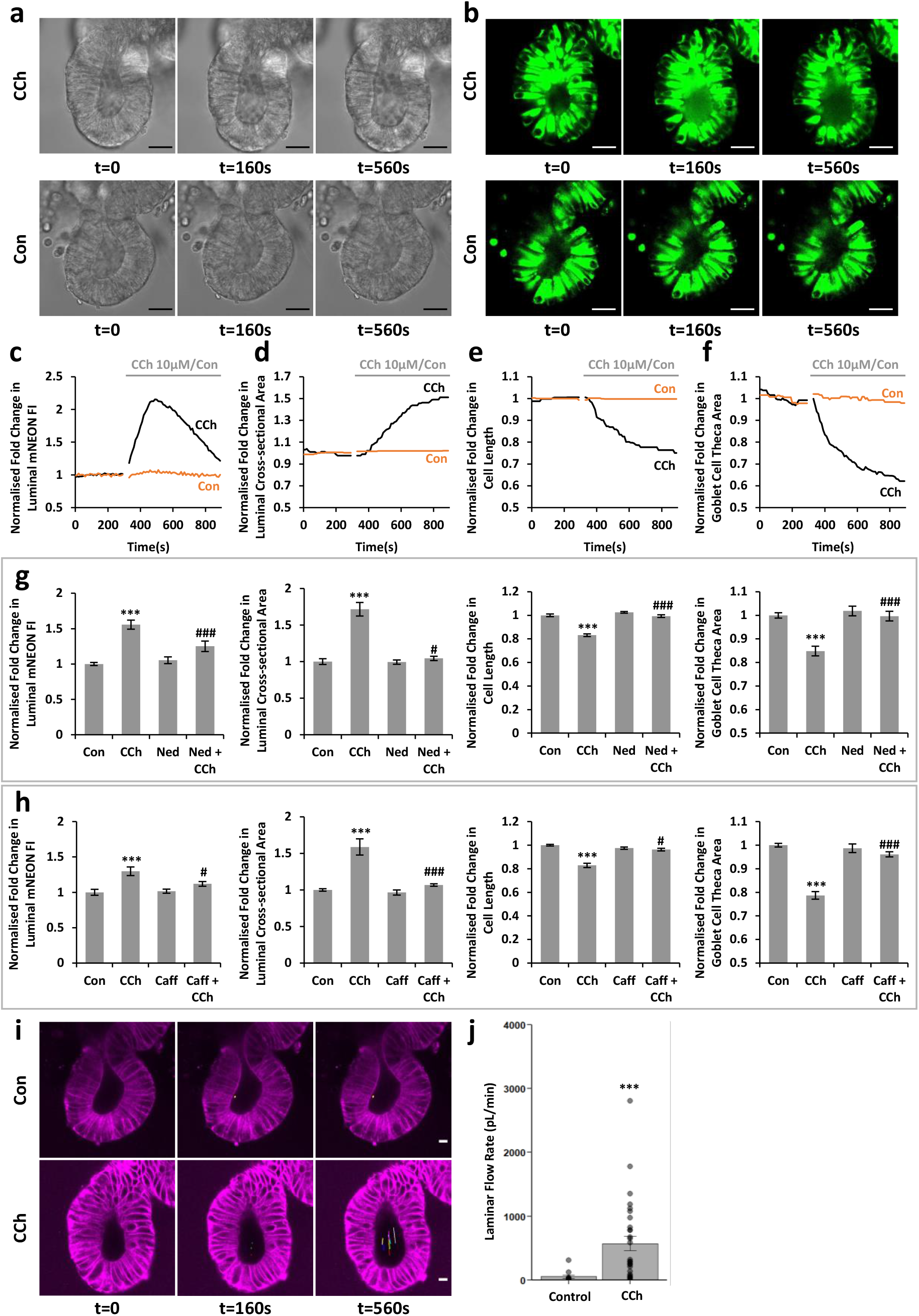
Cholinergic-induced mucus secretion into the crypt lumen is dependent on calcium mobilisation and is accompanied by luminal flushing of the stem cell niche. Representative bright field (a) and MUC2-mNEON (b) time course images of MUC2::mNEON organoid crypt-bases in the presence and absence of CCh (10 µM) stimulation. (c) Kinetics of relative luminal MUC2-mNEON fluorescence intensity, (d) luminal lumen morphometry, (e) apico-basal dimension of crypt base cells and (f) goblet cell theca area in the presence and absence of CCh (10 µM) stimulation. Effects of intracellular calcium channel blockers (g) Ned-19 (500 µM) and (h) caffeine (10 mM) on CCh (10 µM)-induced changes in luminal MUC2-mNEON fluorescence, crypt lumen morphometry, apico-basal dimension of crypt base cells, and goblet cell theca area. (i) Matching deep red fluorescence images of the same organoid crypt bases optimised for tracking the position of luminal particulate with respect to time. (j) Computed laminar flow rate based on maximal velocity of luminal particulate migration. Scale bars = 25µm. ***P<0.001 compared to Control; ###P<0.001 compared to CCh, #P<0.05 compared to CCh. For (g) N=2, n>4, for (h) N=4, n>8, for (j) N=15, n=42. Kruskal-Wallis rank sum tests were undergone for (g): cross-sectional area, theca area; (h): cell length, theca area. One-way ANOVAs were undergone for (g): mNEON fluorescence, cell length; (h) mNEON fluorescence, cross-sectional area. See Supplementary Movie 4.

Finally, we sought to determine the fate of mucus expelled into the stem cell niche. Luminal flow was visualised by pre-incubating MUC2-mNEON organoids with a plasma membrane fluorescent dye (Fig. 7i). On stimulation with CCh the number of fluorescent particles in the lumen increased and their position within the lumen was tracked with respect to time (Fig. 7i; Supplementary Movie 4). Luminal particulate migrated in a collective stream away from the crypt-base towards the opening of the lumen (Fig. 7i; Supplementary Movie 4). The maximum particle velocity in the central lumen was used to compute the laminar flow rate (Fig. 7j)^16^. By contrast, under control conditions, the luminal particulate was minimal and relatively static. To assess a contribution of fluid secretion to cholinergic-stimulated luminal flushing, closed organoids were selected for organoid swelling assays^37^. CCh stimulation increased organoid luminal cross-sectional area over a period of 2 hours; this was inhibited by Clotrimazole, a blocker of calcium-dependent K^+^ channels and an inhibitor of Cl^-^ secretion, the main driver for fluid secretion in the gut (Supplementary Fig. 8a)^38^. Similarly, suppression of cholinergic calcium signal generation by inhibition of TPCs (with tetrandrine or Ned-19), InsP3Rs (with caffeine), or RyRs (with dantrolene) also abrogated CCh-induced organoid swelling, an established proxy for intestinal fluid secretion (Fig. 8 and supplementary Fig. 8)^37^.

**Figure 8.**
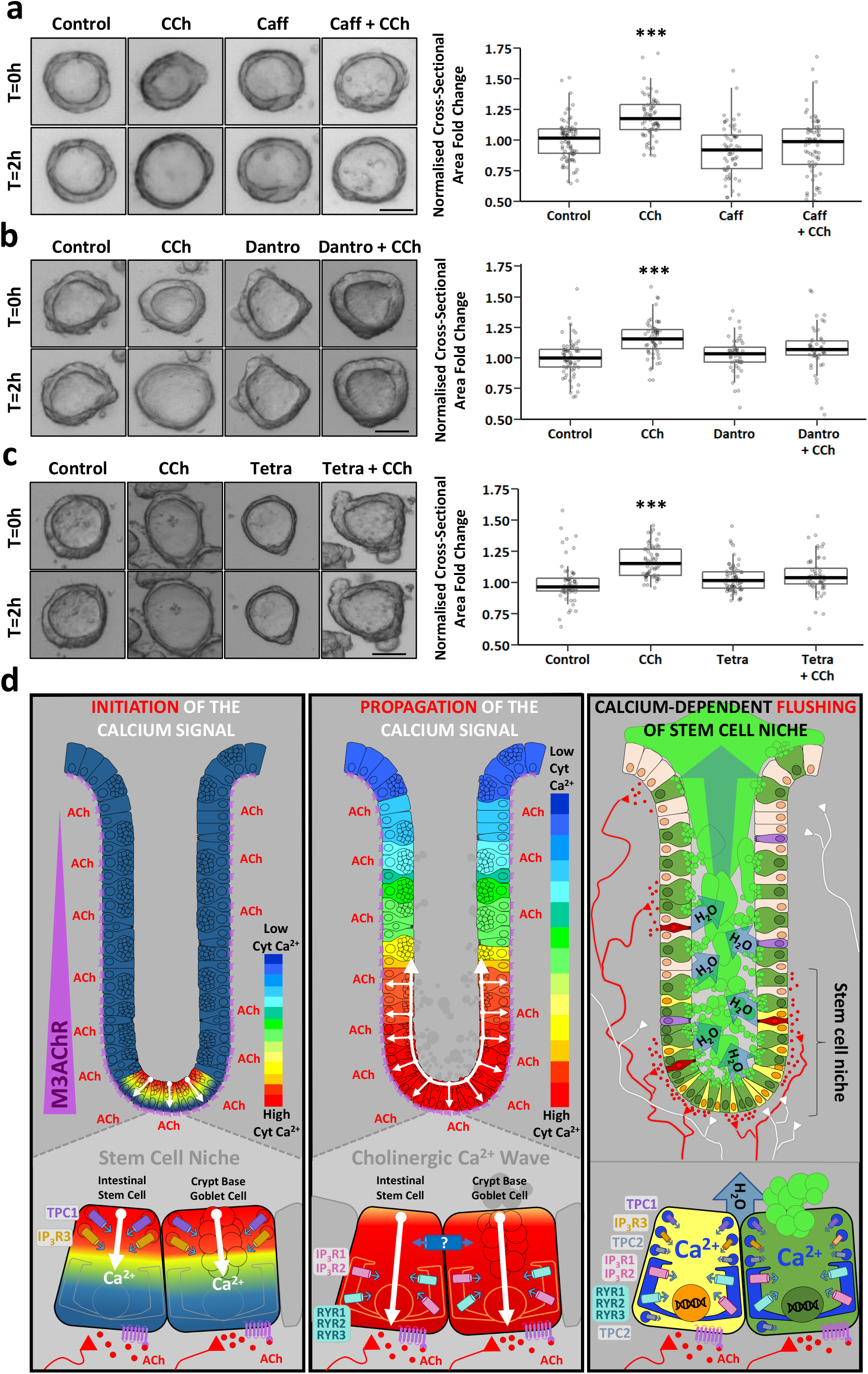
Cholinergic calcium signals coordinate secretion of a fluid rich mucus to flush the human colonic crypt stem cell niche. Matching bright field images and box-plots of Carbachol-stimulated (10 µM, 2 hrs) organoid swelling assays in the presence and absence of treatment with (concentration, pre-treatment): (a) InsP3R inhibitor Caffeine (10 mM, 30 min.), (b) RYR inhibitor dantrolene (50 µM, 2 hrs) or (c) TPC inhibitor Tetrandrine (20 µM, 2 hours). (d) Schematic illustration of calcium-dependent cholinergic excitation-mucus/fluid secretion coupling in the human colonic stem cell niche: left hand panel – cholinergic calcium signal initiation in trigger zone at apical pole of ISCs by TPC1-InsP3R3 cross-coupling; middle panel –activation of MUC2 granule exocytosis, apical-to basal propagation of intracellular calcium wave via InsPR1/2 and RyR1/2/3, and propagation of colonic crypt calcium wave with possible involvement of intercellular communication; Right hand panel – cholinergic calcium signal activation of hydrated mucus secretion to flush the luminal contents of the human colonic stem cell niche. Scale bar = 50 µm. *** P<0.001, significantly different to all other groups. P values calculated via a Kruskal Wallis rank sum test with a Dunn post-hoc. n≥40 organoids; N ≥2 independent experiments in each group.

## DISCUSSION

A major challenge in the field of human gut epithelial biology is to understand the signalling pathways that coordinate diverse physiological processes to maintain intestinal tissue homeostasis. Here we report the discovery of cholinergic calcium signals that couple the secretion of mucus and fluid to flush luminal contents from the microenvironment of the stem cell niche. Importantly, these outcomes have been derived from a combination of human tissue model systems that preserve tissue topology, polarity and cellular diversity, and from complementary use of reporters that enable the mechanism of calcium signals and cellular secretion to be investigated in space and time.

The concept of a cholinergic niche at the base of human intestinal crypts (see Fig. 1i for schema) was envisaged more than 20 years ago with the discovery that cholinergic input stimulated calcium signals at the base of mouse small intestinal crypts^39^ and rat colonic crypts^22^. This was subsequently confirmed in the human colonic epithelium^9, 25^. Our current finding that neuronal and non-neuronal sources of acetylcholine (i.e. cholinergic neurons and cholinergic tuft cells) are concentrated at the human colonic crypt base is consistent with recent observations made in the murine gastric mucosa^19^ and small intestine^20, 40^. We also show that human colonic epithelium M3AChR expression predominates in stem cells and goblet cells in the crypt base. Similarly, M3AChRs have been demonstrated to be expressed in mouse small intestinal Paneth cells and stem cells at the crypt base^20, 41^. In accordance with expression by intestinal stem cells, a role for M3AChRs in mucosal regeneration^20, 40^ and gastrointestinal tumourigenesis^19, 42, 43^ has been proposed. Taken together with our current study, these observations suggest that the cholinergic niche simultaneously regulates both stem cell-driven tissue renewal and antimicrobial defence by secretion of mucus and fluid, the two main contributors to intestinal barrier function. The current study highlights a long suspected fundamental role for calcium in coupling these secretory events and moreover provides a long sort after mechanism for signal generation.

M3AChRs are coupled to cholinergic calcium signals that emanate from a sub-apical microdomain in ISCs and crypt-base-GCs. Juxtaposition of endosomes positive for TPC1 and InsP3R3 positive puncta and the pharmacological profile of intracellular calcium channel inhibitors favour a cooperative cross coupling model for the cholinergic calcium trigger zone^44^. Caffeine was the most potent InsP3R inhibitor of cholinergic calcium signals and has been shown block InsP3R3^29^. Xestospongin C and 2-APB were without effect on the cholinergic response but do block purinergic calcium responses in human colonic crypts (Lee and Williams, unpublished observations). Subapical expression of InsP3R3 has also been demonstrated in cholangiocytes and implicated in bile secretion in health and disease^45^. TPC1 was present in the calcium trigger zone in the intestinal stem cell niche and was accompanied by expression of CD38, which has been implicated in modulation of NAADP levels^26^, an activator of TPCs^46^. The expression of TPC1 in acidic Rab11 recycling endosomes has been documented previously^47^. It is not yet clear whether TPC1 endowed acidic calcium stores prime the neighbouring InsP3R3-positive (ER) calcium stores or vice versa. There is much interest in calcium signalling microdomains and membrane contact sites between different organelles as a means of calcium signal propagation and coordinating inter-organelle cell biology and metabolism^48^ . Given the limited diffusion coefficient of cytosolic calcium, and according to our immunolocalisation studies, it would appear that globalisation of the calcium signal from the apical pole to the basal pole of cells within the intestinal stem cell niche is mediated by regenerative calcium-induced-calcium release by InsP3R1,2 and/or RYR1,2,3; TPC2 may also play a role. Accordingly, in the current study, RYR channel blockers suppressed the amplitude of the cholinergic calcium response. The relative contribution of these calcium channel subtypes is under investigation, as is the wider physiological significance.

The cholinergic calcium signal trigger zone in the apical pole of crypt-base goblet cells is located at the expected site of mucus granule exocytosis^21^. Complementary MUC2 immunolabelling studies and real-time imaging of MUC2-mNEON secretion demonstrated that cholinergic calcium signals promoted calcium-dependent exocytosis of MUC2 granules at the apical membrane of crypt-base-goblet-cells in the human colonic stem cell niche (see Fig.8d for schema). Exocytosis of secretory granules has previously been described as being restricted by the actin cytoskeleton but can be liberated by oscillations in cytosolic calcium^49^. Elevated calcium oscillations have been shown to promote dissociation of calcium sensors such as KChIP3 from MUC5AC granules thereby triggering calcium-dependent synaptotagmin-driven formation of a SNARE complex and granule fusion to the plasma membrane^50–52^. Acidic endolysosomal calcium stores have been implicated in redox stimulated, calcium-mediated mucus granule exocytosis^17^, although the nature of the intracellular calcium release channels was not investigated. It is likely that aspects of the calcium signalosome described by this study of the stem cell niche will be applicable to regulation of calcium-dependent mucus exocytosis from differentiated cryptal and intercryptal goblet cells located at the crypt opening and surface of the gut^6, 13, 53^. Interestingly, zymogen granules in pancreatic acinar cells^54^ and cytolytic T cells^55^ not only display calcium signalling microdomains and membrane contact sites between calcium storage organelles, but the zymogen and cytolytic granules serve as a calcium store that regulates their own exocytosis. This remains to be proven to be the case in intestinal mucus granules.

We also demonstrate that cholinergic calcium signals couple mucus secretion to fluid secretion. Cholinergic-induced fluid secretion is highly regulated by calcium and transient in nature, probably to flush the crypt lumen with minimal fluid loss^9, 56^. Not surprisingly, organoid swelling assays exhibited a moderate size increase. The molecular machinery for calcium dependent chloride-driven fluid secretion is expressed in colonic crypt base cells, including NKCC1^9^ on basolateral membranes and TMEM16A/ANO1^57^ on apical membranes. However, closer inspection of cell type specific labelling of participating ion channels and transporters, and modes of intercellular calcium signalling will be required to confirm whether the cholinergic input stimulates GC mucus granule exocytosis and fluid secretion in the same cell or neighbouring cells, and whether it be in a coordinated or independent manner. In conclusion, this study demonstrates for the first time that input from a cholinergic niche elicits calcium signals in ISCs and GCs that are coupled to the secretion of mucus and fluid which flushes the luminal microenvironment of the colonic stem cell niche. The platform utilised in this current study provides a tractable system to investigate the mechanisms by which calcium signals couple to mucus granule exocytosis and fluid secretion, and to study the status and function of this pathway in human health and disease.

## MATERIAL AND METHODS

### Human colorectal tissue samples

This research study was performed in accordance with a favourable ethical opinion by the Faculty of Medicine and Health Sciences Research Ethics Committee (University of East Anglia; ref. no. 013/2014 – 62 HT). Governance of human tissue procurement for this research project was overseen by the Norwich Research Park Biorepository (IRAS No. 130478). Tissue samples were obtained by consultant histopathologists from the healthy mucosa of patients undergoing surgical resection for colorectal cancer.

### Human colonic crypt isolation and culture

Colonic crypts were isolated as described previously^8, 9^. In short, fresh mucosal tissue samples were collected in ice cold PBS, transported to the laboratory, and incubated in Hepes-buffered saline (HBS): (mM) NaCl 140, KCl 5, Hepes (N-2-hydroxyethylpiperazine-N2-ethanesulphonic acid) 10, d-glucose 5.5, Na_2_HPO_4_ 1, MgCl_2_ 0.5, CaCl_2_ 1, and placed in HBS, which was devoid of both Ca^2+^ and Mg^2+^, and supplemented with EDTA (diaminoethanetetraacetic acid disodium salt) (1mM), for 1 h at room temperature. Crypts were liberated by serial rounds of vigorous shaking, crypt sedimentation and collection. Sedimented crypts were collected and mixed in Matrigel and a 20 µl droplet containing 50-100 crypts was placed onto no. 0 glass coverslips (VWR) contained within a 12 well plate. After polymerisation at 37°C for 5-10 mins, crypts were flooded with 0.5 mls of human colonic crypt culture medium (hCCCM), a variant of that described previously for human intestinal stem cell/organoid culture^58^: advanced F12/DMEM containing B27, N2, n-acetylcysteine (1 mM), Hepes (10 mM), penicillin/ streptomycin (100 U/ml), L-Glutamine (2mM), Wnt-3A (100 ng/ml), IGF-1 (50 ng/ml), Noggin (100 ng/ml) or Gremlin-1 (200 ng/ml), RSPO-1 (500 ng/ml), and the ALK 4/5/7 inhibitor A83-01 (0.5 µM). hCCCM was changed every two days and was modified further according to the stated experimental conditions.

### Human organoid culture

Human colonic organoids were obtained after the passage of colonic crypts grown for 7 days. The samples were detached from the bottom of the plate by manually breaking the Matrigel-containing crypts and these were mechanically dissociated into smaller fragments using a pipette. The suspension containing the crypt fragments was transferred into centrifuge tubes and pelleted at 4°C. Following the removal of the supernatant containing cell debris, crypt fragments were resuspended in hCCCM and the final pellet was embedded in Matrigel, left to polymerise, and the organoids were flooded with hCCCM. Organoids were cultured at 37°C and 5% humidified CO2, fed every 3 days and passaged every 5 to 7 days.

### Generation of single cells

Single intestinal epithelial cells were generated from crypt or organoid culture. Briefly, organoid fragments collected from 3 wells of a 24 well plate were incubated with TrypLE express cell dissociation enzyme (ThermoFisher) supplemented with Y-27632 (10 µM, Sigma) for 15 minutes at 37°C with frequent pipetting. Single cells were washed 3 times and filtered through a 20 µm filter before seeding in Matrigel, left to polymerise and flooded with hCCCM containing Y-27632. Single cells were cultured at 37°C in 5% humidified CO2.

### Whole mount immunohistochemistry

On day 1 of culture following embedding in Matrigel, cultured-crypts, organoids or single cells were fixed with 4% PFA for 1 hour or fixed in a methanol:acetic acid solution (3:1) for 5 minutes, and permeabilised with either SDS (1%) or Triton X-100 (0.5% w/v PBS, 30 min). Non-specific binding sites were blocked with 10% donkey serum and 1% bovine serum albumin for 2 h and washed with PBS. Crypts or organoids were incubated with primary antibodies (1:100-200 dilution) overnight at 4°C. Immunolabelling was visualised by using an appropriate combination of species-specific Alexafluor-conjugated secondary antibodies (488, 568, and 647 nm) raised in donkey (Invitrogen). Crypts or organoids were mounted on glass slides with Vectashield containing DAPI (Vector labs) or SYTOX Blue (Invitrogen).

For the processing of native human fixed sections, a small fraction of the colorectal tissue sample was immediately fixed with 4% PFA for 2 hours at ice cold temperature after the surgical resection. The samples were embedded in OCT compound and frozen on liquid nitrogen in isopentane. Sections of 8 µm thickness were cut using a cryostat and processed for immunohistochemistry as described previously.

### Goblet cell depletion assay

Human colonic crypts or organoids on day 1 of culture were pre-incubated at 37°C with relevant pharmacological agents and/or stimulated with CCh (10 μM) prior to fixation in a methanol:acetic acid solution (3:1) for 5 minutes. Samples were processed for immunohistochemistry as described previously and incubated with primary antibodies against MUC2 and E-cadherin. Images were acquired using laser scanning confocal microscopy (Zeiss 510 META or Zeiss 980 Airyscan) with a ×63 (1.4 numerical aperture) lens. Goblet cell mucus depletion was quantified using ImageJ (NIH) by measuring the fluorescence intensity of MUC2-positive goblet cells’ theca of the goblet cells located at the base of the crypt. A minimum of 2 replicates was measured with a minimum of 100 goblet cells quantified per condition.

### Generation of stable genetically modified MUC2-mNEON organoid lines

For the generation of MUC2-mNEON-labelled colonic organoids, the CRISPR-HOT protocol was followed as previously described^23^. In short, the guide RNA targeting exon 49 of the *MUC2* gene was cloned into the pSPgRNA plasmid (Addgene #47108) using the following oligonucleotide sequences: *MUC2* fwd, 5′-CACCGCATCTGGGGAGCGGGTGAGC-3′; *MUC2* rev, 5′-AAACGCTCACCCGCTCCCCAGATGC-3′. Human colonic organoids were collected from 3 wells from a 24 well plate and mechanically dissociated into small fragments. Following dissociation, organoid fragments were washed and resuspended in Opti-MEM supplemented with Y-27632 (10 µM) and 10µg of pSPgRNA plasmid together with the frame selector plasmid pCAS9-mCherry-Frame +1 (Addgene #66940) and the mNEON targeting plasmid (a kind gift from V. Hornung^36^). Transfection was performed using cuvette electroporation using the NEPA21 electroporator (Nepa Gene) with the following parameters: Poring Pulse (Voltage = 175 V, Pulse Length = 7.5 msec, Pulse Interval = 50 msec, Number of Pulse = 2), Transfer Pulse (Voltage = 20 V, Pulse Length = 50 msec, Pulse Interval = 50 msec, Number of Pulse = 5), as previously described^59^. Fragments were supplemented with 1ml of hCCCM supplemented with Y-27632 (10 µM) and allowed to recover for 30mins before seeding in Matrigel and cultured in hCCCM supplemented with Y-27632 (10 µM). Electroporation success was assessed by transient mCherry expression and 2-3 days after electroporation at which time MUC2-mNEON positive organoids were already visible. Organoids expressing fluorescent MUC2-mNEON were picked and selected to establish a clonal knock-in organoid line, and its genotype characterised by PCR amplification followed by Sanger sequencing.

### Confocal microscopy

Following immunohistochemistry, whole-mounted microdissected (ie, native) and cultured crypts were visualised by laser scanning confocal microscopy (Zeiss 510 META or Zeiss 980 Airyscan). A ×63 (1.4 numerical aperture) objective was used to obtain confocal images of the longitudinal crypt-axis. Image stacks were taken at 1–3 μm intervals which allowed selection of precise focal planes. The same acquisition parameters were used prior to post-hoc comparison of immunolabelling fluorescence intensity. Image analysis was performed using ImageJ. Three dimensional images were rendered in Volocity (Improvision).

### RT-PCR

RNA was extracted from freshly isolated colonic primary mucosa, freshly isolated crypts and from human colonic organoids after their 4^th^ passage, and isolated using the ReliaPrep™ RNA Miniprep System (Promega) according to the manufacturer’s instructions. Complementary DNA was generated using 500 ng of total RNA, oligo-dT primers (Promega) and M-MLV Reverse Transcriptase kit (Thermo Fisher Scientific). RT-PCR was performed using the G-Storm thermal cycler on 25 µl reaction samples containing forward and reverse primers (200 nM) listed in Table 1, dNTPs (200 µM, Promega), 0.04 U/μl of GOTaq® G2 DNA Polymerase (Promega), PCR buffer (Promega) and MgCl2 (2.5 mM). RT-PCR products were run on a 2.5% agarose gel and visualised by ethidium bromide staining.

For the analysis of the knock-in efficiency and genotyping of MUC2-mNEON organoids, genomic DNA was isolated using ReliaPrep™ gDNA Tissue Miniprep System (Promega) according to the manufacturer’s instructions and was then amplified using 0.02 U/ μl of Q5® High-Fidelity DNA Polymerase (NEB) according to manufacturer’s instructions with the primer sets listed in Supplementary Table 1.

### Calcium imaging

Isolated colonic crypts or organoids were loaded at RT with Fura-2/AM (5 μm, 2h; Thermo Fisher Scientific) in HBS, to monitor Ca^2+^ as previously described^9^. The loaded specimen was placed in an experimental chamber located on the stage of an inverted epifluorescence microscope (Nikon TE200) using a ×40 1.1 NA objective. Experimental solutions were administered via a two-way tap. The activity of the correspondent calcium channels was blocked using the following inhibitors: Caffeine (10 mM, Sigma), Trans-Ned-19 (500 µM, Tocris), Dantrolene (30 or 50 µM, Tocris), Ryanodine (50 µM, Tocris), Tetrandrine (20 µM, Sigma), Diltiazem (250 µM, Tocris), CPA (20 µM, Tocris), 1,1-Dimethyl-4-diphenylacetoxypiperidinium iodide (4-DAMP, 0.1 µM, Tocris), U73122 (10 µM, Sigma) and Bafilomycin A1 (2.5 µM, Tocris). Fluorescence ratio data are presented in the form of pseudocolour images or as traces, where the average ratio values within regions of interest (ROIs) located at the base of the crypt are plotted with respect to time.

For the analysis of calcium signal initiation at the cellular level, colonic organoids were loaded at RT with Fluo-8/AM (5 μM, 2h; Abcam) or Calbryte™ 630 AM (10 μM, 2h; AAT Bioquest) in HBS. To aid cellular loading, the working solution also contained 0.04% Pluronic® F127 (Sigma) and Probenecid (2.5 mM; AAT Bioquest). Samples were imaged using super-resolution confocal microscopy using a Zeiss 980 Airyscan microscope equipped with a x40 1.3 NA objective. Images were noise subtracted using the average of the first 10 frames prior to addition of CCh using ImageJ’s Times series analyser plug in and the changes in fluorescence intensity are presented as pseudocolour images.

### Organoid swelling assay

Human colonic organoids embedded in Matrigel were pre-incubated on day 1 with hCCCM containing the relevant pharmacological agents at 37°C and then stimulated with CCh (10 µM). The whole well containing organoids was live imaged for 2 hours after stimulation, using an inverted microscope (Nikon-Ti, 4x objective). The data was analysed using ImageJ through measurement of the diameter of the intraluminal membrane of the organoid (n≥15) at t=0h and t=2h and obtaining the swelling ratio.

### MUC2-mNEON data analysis

MUC2::mNEON organoids were loaded with CellMask™ Deep Red Dye (Thermo Fisher) for 1h (5µg/ml) and pre-incubated with Caffeine (10 mM) or Ned-19 (500 µM) at 37°C prior to stimulation with CCh (10 µM) for 15 minutes. The organoids were imaged at 10 second intervals for 5 minutes prior to stimulation and 15 minutes post stimulation using a Zeiss 980 confocal microscope (x40 oil lens) at 37°C and 5% CO2. Changes in luminal MUC2-mNEON were analysed by measuring the mNEON luminal fluorescence intensity over time. Luminal morphometry was assessed by quantifying the changes in cross-sectional area over time. Secretory volume decrease of crypt base goblet cells was analysed by measuring the cell length of 3 MUC2-mNEON expressing cells at t=0 and t=15 min. Changes in goblet cell theca area were quantified by drawing regions of interest around the theca of the MUC2-mNEON^+^ goblet cells at t=0 and t=15 min, while the kinetic values were obtained by measuring the changes in theca area over the course of the experiment. The data was analysed using ImageJ. Luminal flushing was quantified through tracking of individual deep red positive particles in the lumen of crypt domains and the fluid flow rate for luminal particulate was determined via the following equation derived from Halm and Halm (2000)^16^: 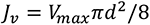. Here, Jv= total volume flow, Vmax=velocity of the fastest moving luminal particle, d= lumen diameter.

### Statistical analysis

Data is expressed as means ± SEM (n is the number of crypts/organoids derived from N patients). Comparisons between two groups was performed using either an unpaired t-test or Mann Whitney U test where appropriate. Pairwise comparisons between >2 conditions were analysed using a one-way ANOVA alongside a Tukey post hoc, or a Kruskal-Wallis rank sum test alongside a Dunn post hoc with a Bonferroni correction where appropriate. The number of human tissue donors, number of crypts or organoids, the number of replicates and independent experiments, as well as the type of statistical analyses performed, is indicated in the figure legends.

## Acknowledgements

We thank Roxanne Brunton-Sim and Rachael Stanley (Norwich Research Park Biorepository) for their help with human tissue research governance. Staff at Same Day Admissions Unit, Norfolk and Norwich University Hospital were extremely helpful when taking patient consent. Paul Thomas (BIO imaging platform, UEA) is thanked for his expert guidance on imaging modalities. Prof. Dr. Veit Hornung kindly gifted the mNEON plasmid for generation of MUC2::mNEON human organoid line. This paper is dedicated to the memory of Mr Les Rhoades, Chairman of Humane Research Trust; Les, your legacy lives on.

## Author contribution

Experimental design (MW, VJ, NP-L, AP, NJ); Manuscript writing (MW, NP-L, VJ); figure preparation (MW, NP-L, VJ, AL, ST); image processing and analysis (NP-L, VJ, CK, ST, JC, MW); colonic crypt culture and organoid culture (MW, VJ, AP, NP-L); mucus depletion assay (VJ, NP-L, CK, AL); RT-PCR (NP-L), MUC2::mNEON organoid generation (NP-L, VJ, BM); immunofluorescent labelling (VJ, NP-L, AP, AL, CK, JC); calcium imaging (AL, NP-L, ML, GR); Provision of clinical tissue samples and tissue dissection (JH, SK, CS, AT, AS, AP, IS, DB, AP, VJ, MW).

---

**Supplementary Figure 1.**
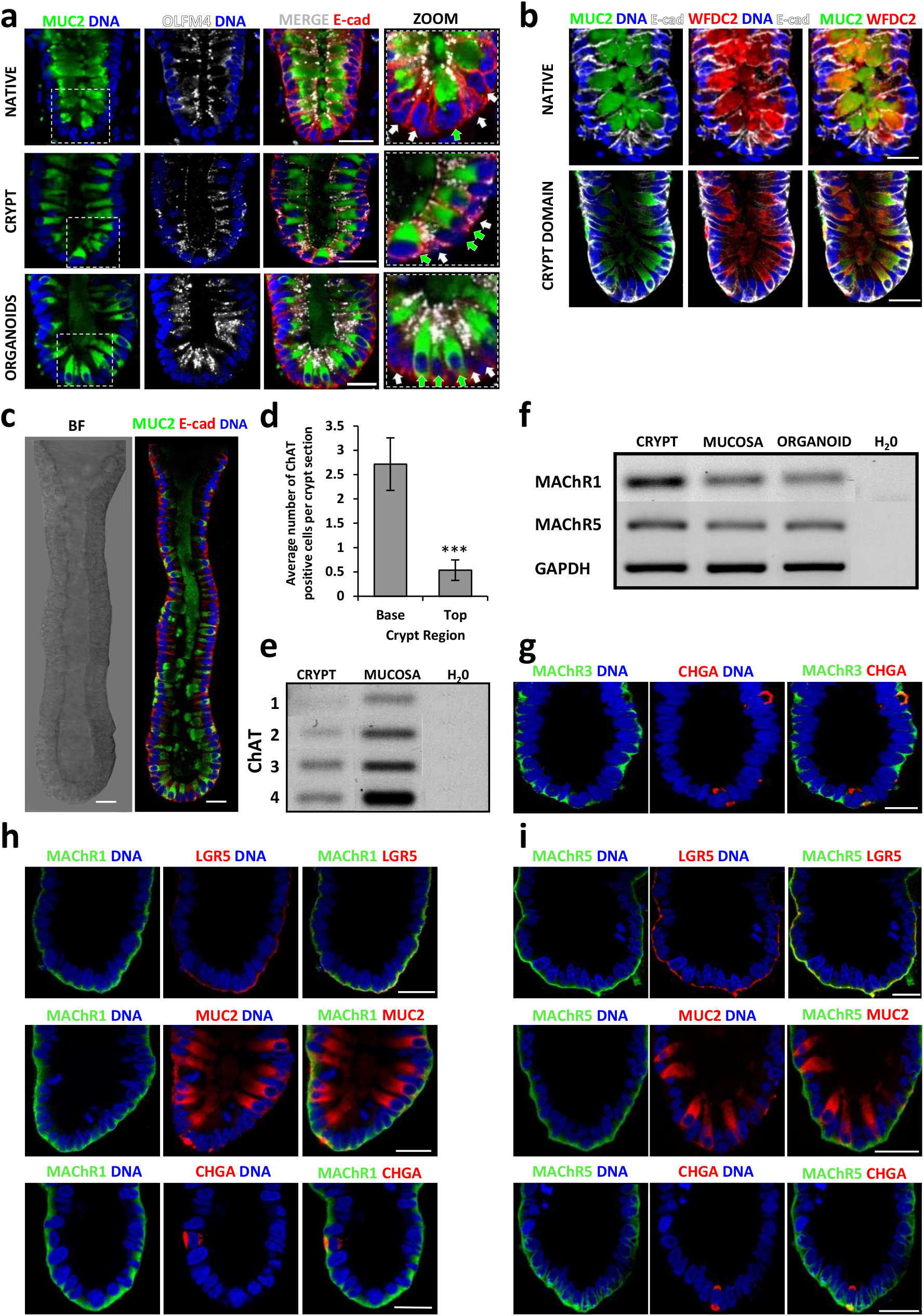
Cellular composition of the cholinergic-stem cell niche at the base of human colonic crypts and organoids. (a) Immunofluorescent labelling of OLFM4 (white)-positive stem cells sandwiched between MUC2 (green)-positive goblet cells at the base of native human colonic crypts, cultured colonic crypts and budding colonic organoids. (b) Immunofluorescent co-labelling of crypt-base goblet cells with MUC2 (green) and WFDC2 (red) in native and cultured crypts. (c) Series of stitched images representing immunofluorescent labelling of MUC2 (green) goblet cells along the cultured colonic crypt-axis; BF - bright field, E-Cad – e-cadherin. (d) Quantification of ChAT-positive tuft cells along the native colonic crypt-axis; N=5 subjects; n=28 crypts. (e) RT-PCR detection of ChAT mRNA in native mucosa and isolated colonic crypts. (f) mRNA expression of MAChR1 and MAChR5 in native mucosa, isolated crypts and colonic organoids. (g-i) Immunofluorescence co-labelling of M1, M3 and M5 (green) muscarinic receptor subtypes with markers for intestinal epithelial cell types (Lgr5 – stem cells, MUC-2 – goblet cells, CHGA – enteroendocrine cells; all red). Scale bars 20µm, ***P<0.001. P value calculated via an unpaired t-test.

**Supplementary Figure 2:**
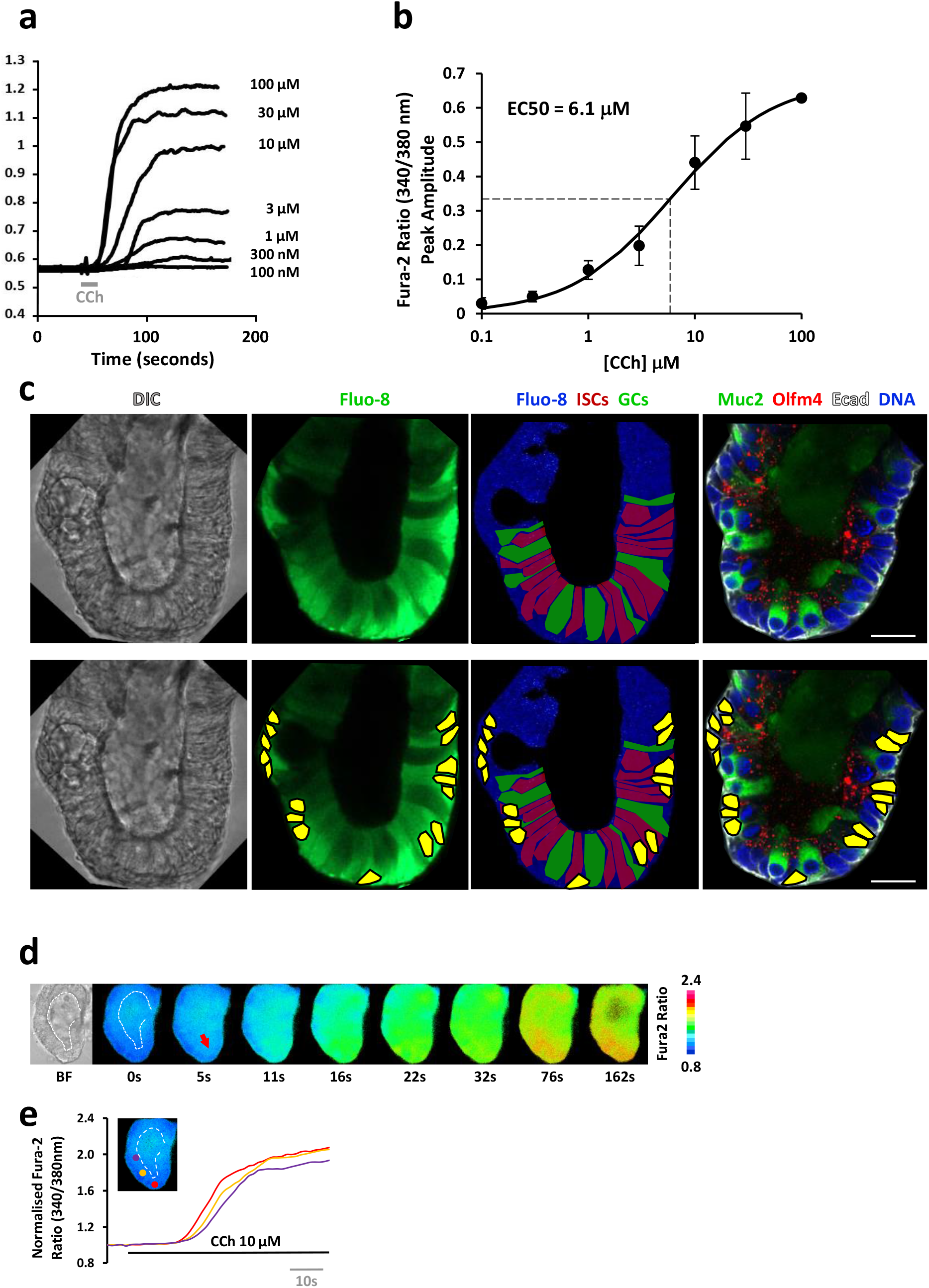
Human colonic organoids recapitulate the characteristics of cholinergic calcium signals associated with the native colonic crypt stem cell niche. (a) Average peak amplitude of different concentrations of CCh used to calculate the EC50 (b) in the colonic stem cell niche; crypts from N=4 subjects, n=3 crypts per data point. (c) Post-hoc immunophenotyping of colonic crypt cell types exhibiting Carbachol-stimulated cholinergic calcium signals (goblet cells - MUC2, green; intestinal stem cells - OLFM4, red). Bottom row highlights nuclei (yellow) that are present in the same focal plane in the live and IF images to aid the immunophenotyping. (d) Fura-2 ratio images of a human colonic organoid stimulated with Carbachol (10 µM) (red arrowhead indicates site of calcium signal initiation) and (b) kinetics of cholinergic calcium signal along the organoid crypt-axis (N>3 subjects, n>50 organoids). Scale bars = 25 µm.

**Supplementary Figure 3:**
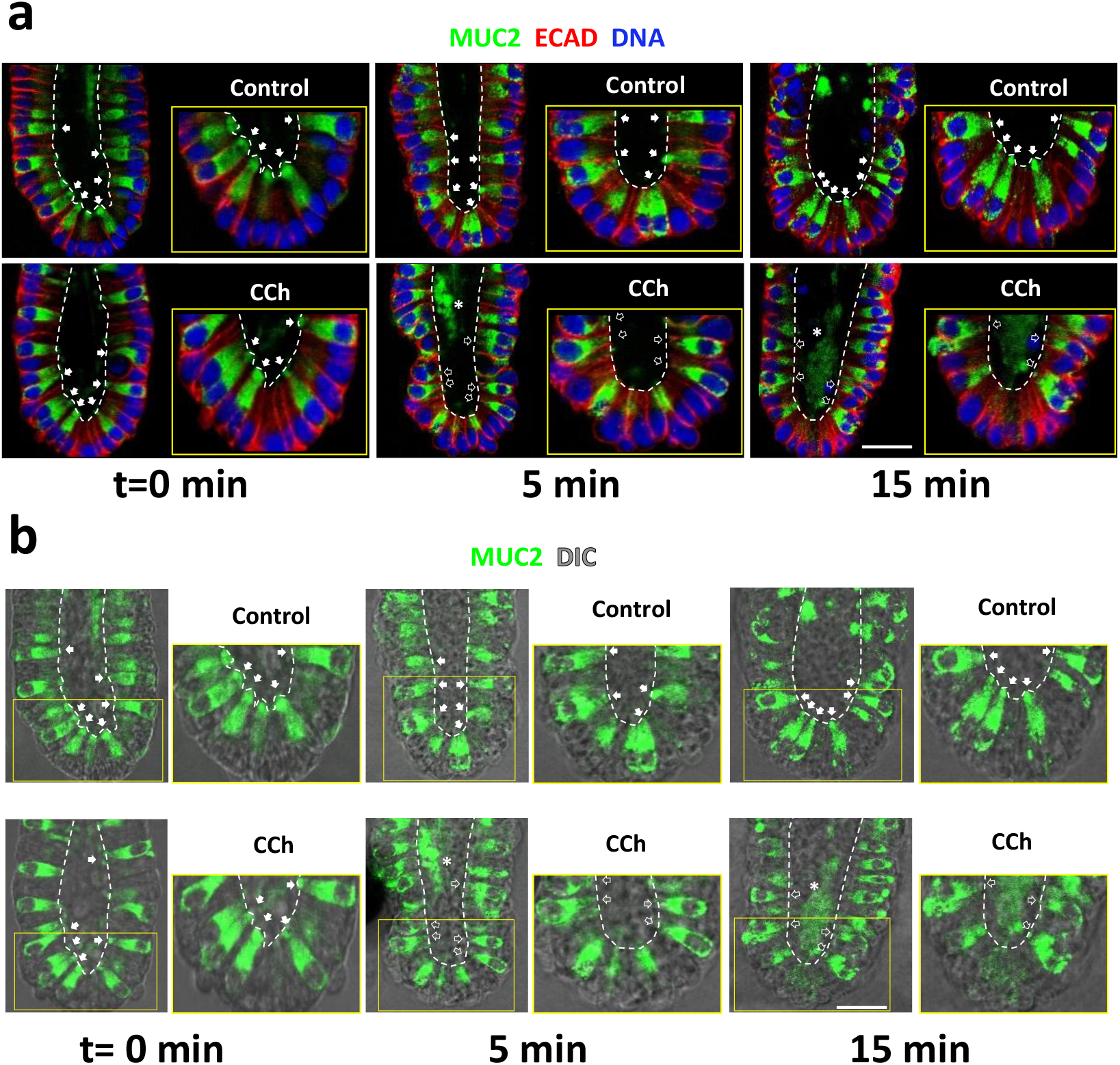
Muscarinergic mobilisation of intracellular calcium induces mucus secretion at the human colonic crypt base. (a) Representative immunofluorescent MUC2 labelling (green) of matched time course for unstimulated controls and Carbachol-stimulated depletion of MUC2 from crypt-base goblet cells; (b) representative merged MUC2 (green) and DIC images of time course for control and Carbachol-stimulated depletion of MUC2 from crypt-base goblet cells; filled arrowheads – goblet cells replete with mucus, empty arrowheads - goblet cells depleted of MUC2, grey filled arrowheads – partially emptied/re-filled MUC2 goblet cells. N=3, crypt≥10, GC>100. Scale bar = 25 µm.

**Supplementary Figure 4:**
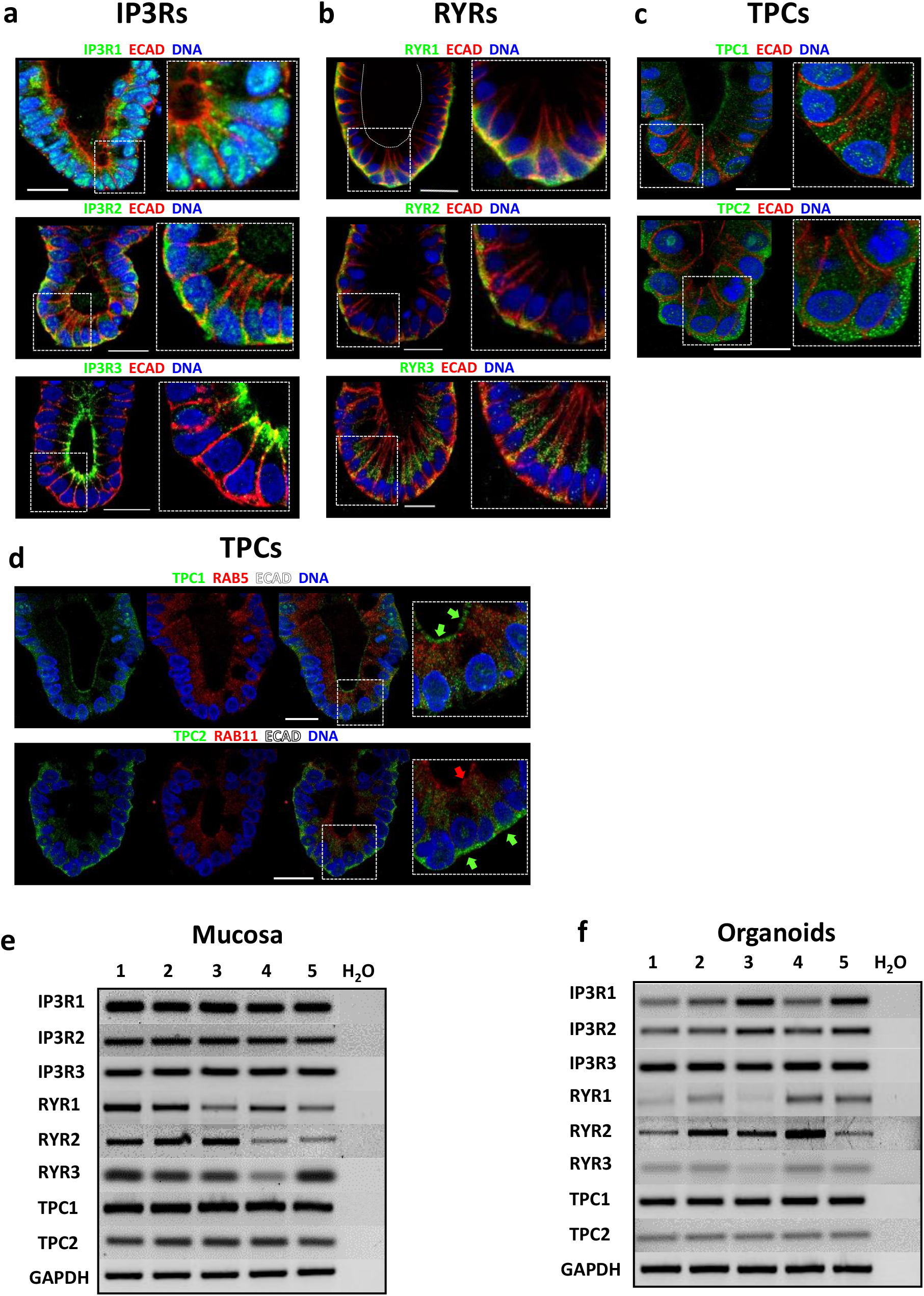
Intracellular calcium channel expression in the native human colonic mucosa and organoids. Immunolocalisation of InsP3R1, InsP3R2, InsP3R3. (a), RYR1, RYR2, RYR3 (b), and TPC1 and TPC2 (c), in combination with E-Cadherin; N ≥ 3 organoid lines, n ≥ 10 organoids; Scale bar = 25µm. (d) Representative immunofluorescence images of TPC1 in combination with early endosomal marker RAB5 and E-cadherin (Top) and TPC2 in combination with recycling endosomal marker RAB11 and E-cadherin (Bottom); N≥3 subjects, n≥10 crypts. Scale bar = 25µm. RT-PCR of intracellular calcium release channel mRNA transcript expression in native human colonic mucosa (e) and organoids (f) (N=5 subjects).

**Supplementary Figure 5:**
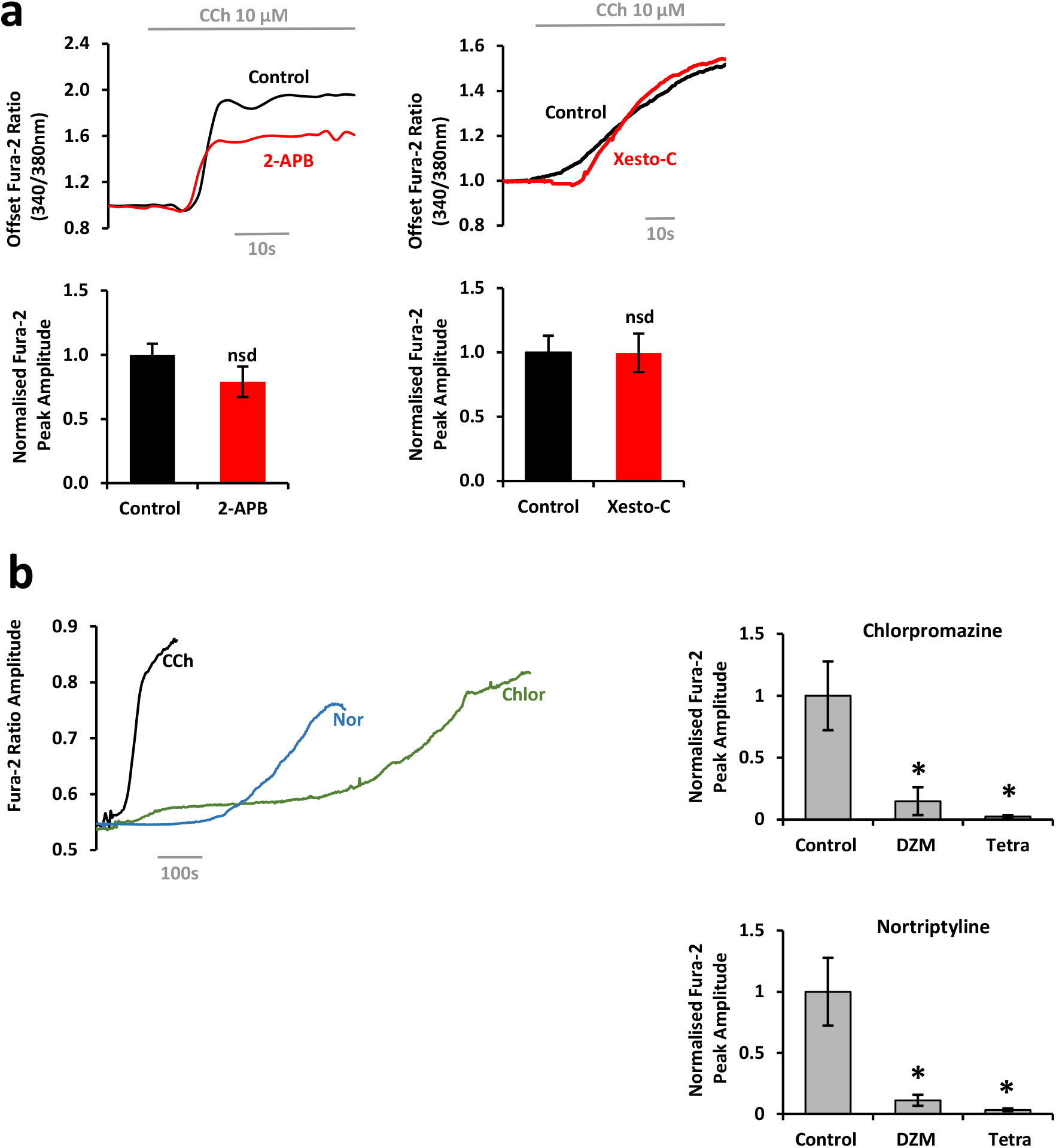
Muscarinergic calcium signals are dependent on cooperation between two pore channels, InsP3 and Ryanodine receptors. (a) Effects of InsP3 receptor inhibitors 2-APB (100 nM, 2 hrs) and Xestospongin-C (4 µM, 2hrs) on Carbachol-induced calcium signal generation and the corresponding bar chart of peak amplitude of the response. (b) Representative Fura-2 ratio traces of CCh (10 µM), Chlorpromazine (100 µM) and Nortriptyline (100 µM) and bar chart of mean peak response amplitude of the effects of two pore channel inhibitors Diltiazem (250 µM, 2 hrs) and Tetrandrine (20 µM, 2hrs) on Chlorpromazine and Nortriptyline calcium signal generation respectively. N≥ 3, crypt≥10in each case. Unpaired t tests were undergone for (a), one-way ANOVAs with a Tukey post hoc was undergone for (b). * indicates P<0.05.

**Supplementary Figure 6.**
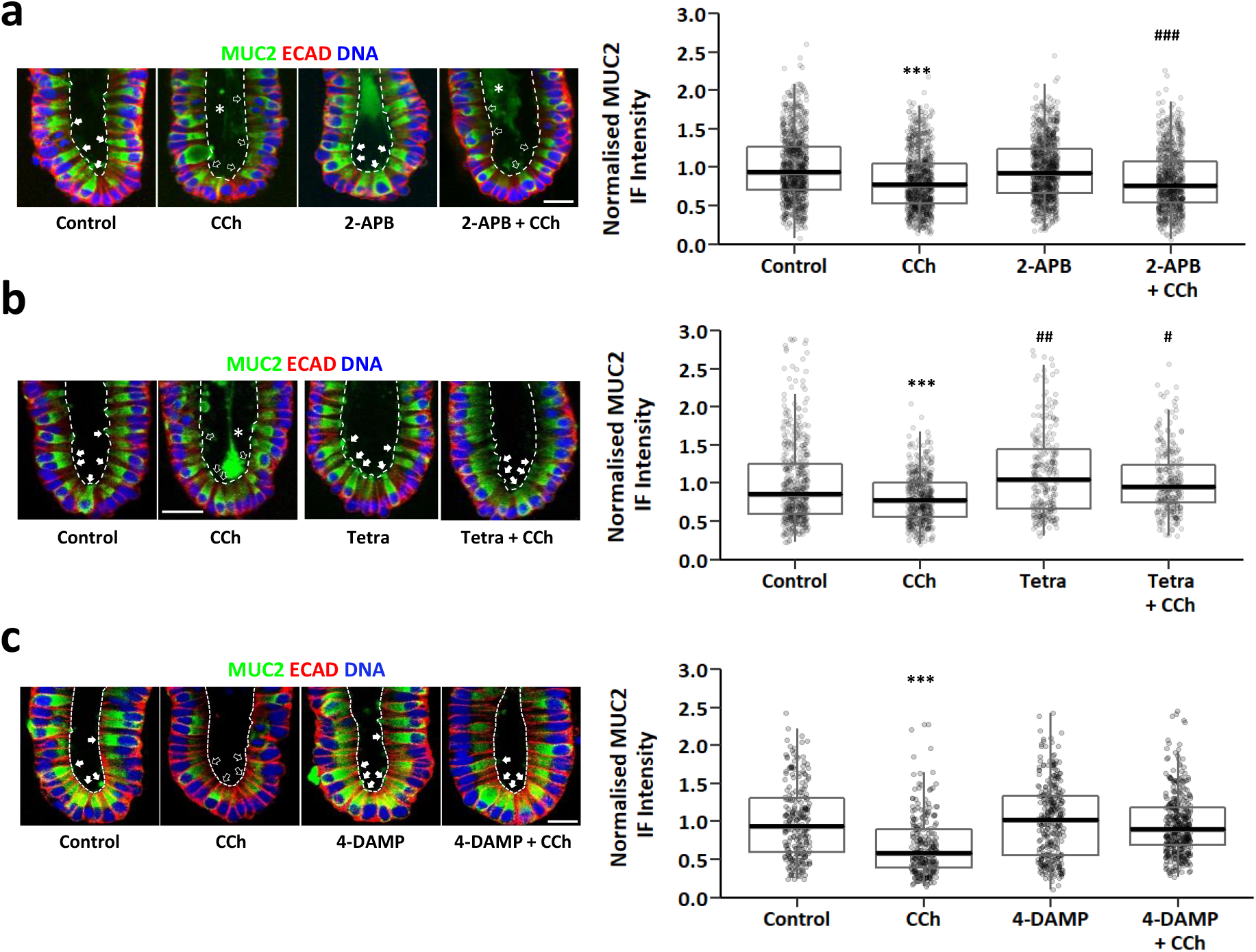
Effects of additional calcium signalling pathway modulators on cholinergic excitation-mucus secretion coupling. MUC2 immunofluorescence images, and accompanying analysis of goblet cell immunofluorescence intensity, following stimulation of human colonic crypts with Carbachol (10 µM, 5 min) in the presence or absence of intracellular calcium channel inhibitors for (a) InsP3Rs (2-APB, 100 µM), (b) TPCs (Tetrandrine, 20µM) and (c) M3AChR inhibitor 4-DAMP (100 nM). N=3, crypt≥10, GC>100 in each case. All scale bars = 25µm. Arrows - secreted mucus granules, empty arrowheads - goblet cells depleted of MUC2, filled arrowheads – goblet cells replete with mucus. ***P<0.001 with respect to all other groups; ###P<0.001 with respect to control; ##P<0.01 with respect to control; #P<0.05 with respect to control. P values calculated via Kruskal Wallis rank sum tests with a Dunn post-hoc.

**Supplementary Figure 7:**
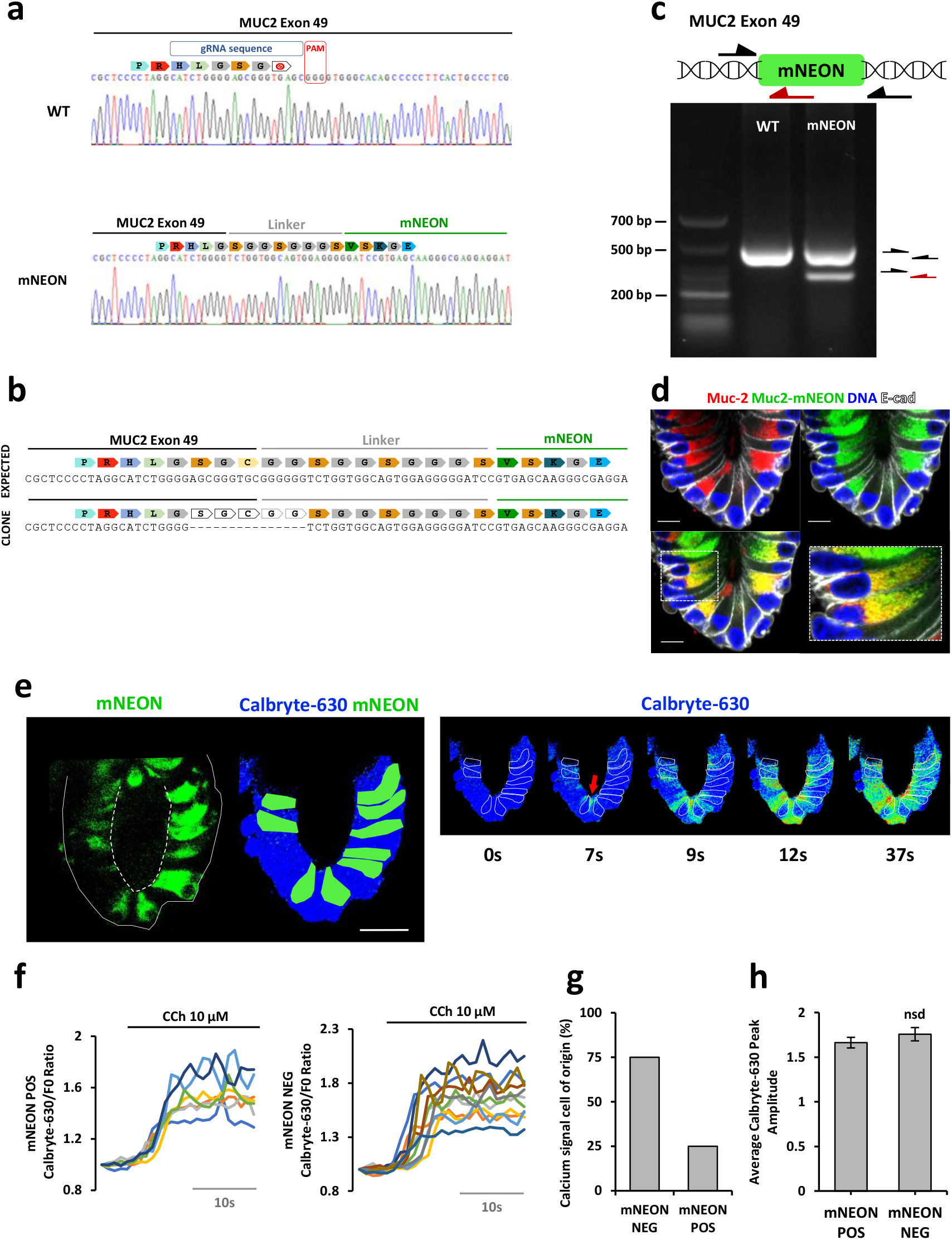
Validation of MUC2::mNEON organoids. (a) Sanger sequencing of WT and MUC2-mNEON genomic DNA confirms in-frame knock-in of the mNEON fluorescent protein at exon 49 of the MUC2 locus. (b) Sequencing results shows the MUC2-mNEON in-frame knock-in clone contains a deletion of 5 amino acids, 3 in exon 49 of the MUC2 gene and 2 in the linker sequence of the knock-in protein, which results in an effective deletion of 1 amino acid from exon 49 (i.e. a cysteine). (c) RT-PCR for genotyping MUC2-mNEON clone using a forward primer located 5’ outside of the mNEON sequence and two reverse primers, one located 3’ outside of the mNEON sequence and another one located inside the mNEON sequence. The resulting two bands on the mNEON lane indicates the presence of both WT and MUC2-mNEON alleles demonstrating heterozygosity of the clone. (d) MUC2-mNEON fluorescence and & MUC2 immunofluorescence demonstrating 100% co-labelling of organoid crypt-base cell types. (e) F/F0 Calbryte-630 fluorescent images of relative intracellular calcium levels in a MUC2::mNEON organoid-crypt-base stimulated by CCh (10 µM); see Supplementary Movie 3. (f) Kinetics of relative cytosolic calcium levels for mNEON^-^ & mNEON^+^ cells. (g) Bar chart for relative frequency of cell-of-origin for cholinergic calcium signal initiation and (h) calcium signal amplitude for mNEON^-^ & mNEON^+^ cells. Calcium data from n≥4 organoids. Scale bars = 25µm.

**Supplementary Figure 8:**
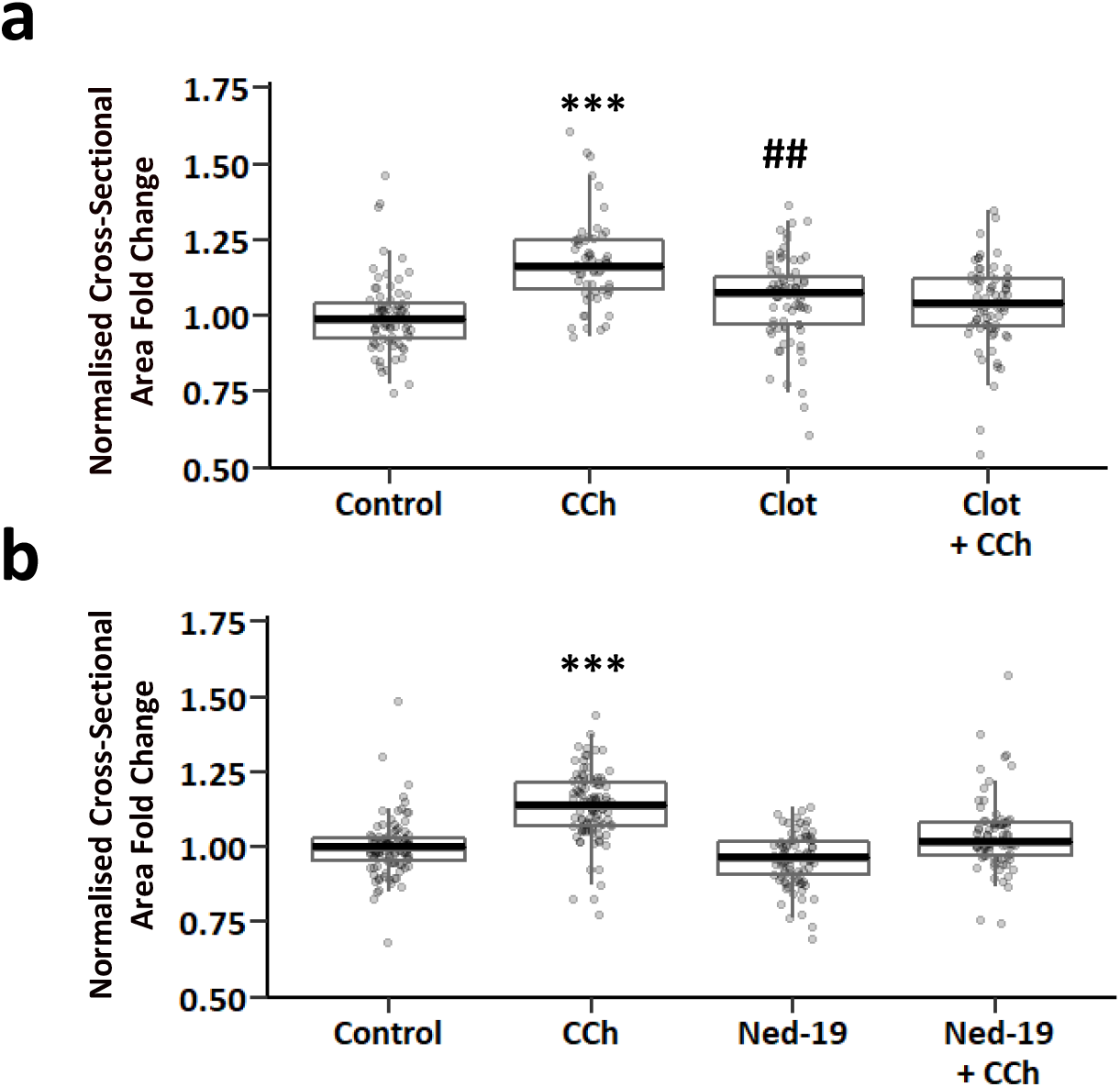
Cholinergic input coordinates excitation-mucus and -fluid secretion coupling to flush the lumen of the human intestinal stem cell niche. Effects of the calcium–sensitive K^+^ channel blocker Clotrimazole and intracellular calcium release channel inhibitor Ned-19 on Carbachol stimulated colonic organoid swelling; summary bar chart of fold-change in organoid luminal cross-sectional area following stimulation with Carbachol (10µM) in the presence of (a) Clotrimazole (50 µM) or (b) Ned-19 (500 µM). N=2 colonic organoid lines, organoids≥20 in each case. ***P<0.001 with respect to all other groups; ##P<0.01 with respect to control. P values calculated via Kruskal Wallis rank sum tests with a Dunn post-hoc.

## SUPPLEMENTARY MOVIES

**Supplementary Movie 1:** Human colonic crypt loaded with Fura-2 and stimulated with Carbachol (10 µM; time = 120s). The cholinergic calcium signal initiates in the stem and propagates up the crypt-axis. Imaging mode = epifluorescence.

**Supplementary Movie 2:** Human colonic crypt loaded with Fluo-8 and stimulated with Carbachol (10 µM; time = 55s). The cholinergic calcium signal initiates in slender intestinal stem cells and propagates throughout the stem cell niche. Imaging mode = confocal fluorescence microscopy.

**Supplementary Movie 3:** MUC2::mNEON crypt-like organoid loaded with Calbryte-630 and stimulated with Carbachol (10 µM; time = 120s). Initial image of MU2-mNEON^+^ goblet cells fades away to reveal a time series of F/F0 images. The cholinergic calcium signal initiates in slender MUC2-mNEON^negative^ cells before registration in MUC2-mNEON^positive^ cells. Imaging mode = confocal fluorescence microscopy.

**Supplementary Movie 4:** MUC2::mNEON crypt-like organoid loaded with Deep Red Cell Mask. Carbachol (10 µM) stimulated the expulsion of MUC2-mNEON into the lumen. The MU2-mNEON theca exhibited a decrease in cross-sectional area whilst the lumen increased in size. Decreasing the threshold on the ‘red’ channel saturated cellular Deep Red fluorescence but revealed flushing of particulate out of the crypt lumen. Imaging mode = confocal fluorescence microscopy.

